# From Red Tides to Healthy Ecosystems by Nutrient Management in Ise Bay and Mikawa Bay

**DOI:** 10.1101/2025.05.13.653706

**Authors:** Ippei Noda, Shinako Iki, Kiyoshi Naruse

**Author notes:** Aichi Prefectural Government, 3-1-2 Sannomaru, Naka-ku, Nagoya, Aichi 460-8501, Japan. Authors: Shinako Iki; Kiyoshi Naruse. **SYNOPSIS.** This study reports that adjusting N and P levels can improve phytoplankton and food chain health without causing harmful algal blooms.

## Abstract

Ise Bay and Mikawa Bay, situated near major urban and industrial areas, have been increasingly affected by water pollution due to human activities. Over time, eutrophication—primarily driven by excess phosphorus (P) and nitrogen (N)—has led to frequent occurrences of red tides and blue tides, significantly impacting coastal fisheries. To mitigate environmental pollution, efforts have focused on purifying the rivers and brackish lakes that flow into these bays. However, as pollution loads have decreased, these bays have transitioned from a eutrophic to an oligotrophic state, leading to a decline in the populations of benthic organisms and their higher consumers. In this study, we examine the relationship between phytoplankton growth—indicated by chlorophyll a (Chl-a)—and nutrient concentrations, particularly P and N. Through this analysis, we determine the optimal levels of P and N necessary to support healthy phytoplankton growth while preventing harmful algal blooms. Based on our findings, we propose a balanced approach to enhancing the nutritional conditions of enclosed coastal waters without triggering red or blue tides.

## INTRODUCTION

Throughout the long history of life on Earth, organisms have evolved to adapt to their environments. Oceans, covering about 70% of the Earth’s surface, are home to a vast array of marine life supported by a food chain based on phytoplankton.(1,2) These phytoplankton not only carry out photosynthesis and absorb carbon dioxide but also produce around half of the Earth’s oxygen, playing a crucial role in maintaining marine biodiversity and the global environment.(3,4) However, economic growth and industrial development had heavily impacted Ise Bay and Mikawa Bay. Due to the proximity of Nagoya and surrounding cities, as well as the concentration of automobile manufacturing and agricultural areas, these bays were suffering from severe pollution, a situation exacerbated by the low fluidity of the seawater within the bay. Pollutants such as domestic wastewater, industrial effluents, and agricultural runoff had deteriorated the water quality, causing frequent red and blue tides and mass deaths of marine life. To restore Ise Bay and Mikawa Bay to their pre-pollution state, efforts have been made to reduce the pollution load flowing into the bays and into the rivers and brackish lake connected to the bays, and to purify the water quality. Sewerage systems were constructed to treat domestic wastewater (Figures **S1c** and **S1e**), significantly lowering the concentrations of organic matter and nutrients.(5) Additionally, dredging and sand-capping were undertaken to improve the bottom sediments of brackish lakes and rivers (Figures **S2a** and **S2b**), leading to gradual improvement in water quality and a decrease in the frequency of harmful algal blooms.(6) Unexpectedly, the reduction of pollutant inflows also caused the bays to become oligotrophic, leading to a decline in fish catches.(7) Pollutant loads such as unregulated ammoniacal and nitrate nitrogen from fertilizers and nitrate nitrogen from wastewater treatment plants further complicate the issue. These pollutants undergo chemical transformations such as denitrification, reducing inorganic nitrogen levels in the waters. Figures **S5a, S5b, S5c**, and **S5d** show that N and P concentrations in Mikawa Bay and Ise Bay decreased significantly from the 1980s to the 2010s, along with a reduction in Chl-a concentrations (Figures **S5e** and **S5f**). Consequently, there was a notable decline in the annual catches of benthic organisms, such as mantis shrimp, conger eels, shrimp, and short-neck clams, as well as higher-order consumers like pufferfish.(8,9) This phenomenon suggests that the decrease in phytoplankton disrupts the food chain, including zooplankton, thereby affects the growth of various marine organisms. However, rapidly increasing levels of P or N to alleviate nutrient deficiency risks the recurrence of harmful algal blooms such as red and blue tides. Recently, it has been reported that the requirements for nitrogen and phosphorus vary depending on the trophic level(10,11), and it has been pointed out that this relative ratio is important in mitigating the adverse effects of oligotrophy on ecosystems. Therefore, haphazard adding inorganic N and inorganic P is not a viable solution. To effectively address nutrient deficiencies, it is important to understand the current nutrient status of Mikawa Bay and Ise Bay, identify the root causes of phytoplankton decline, and minimize nutrient fluctuations. In this study, we investigate the long-term changes in phytoplankton distribution in Ise Bay and Mikawa Bay, and propose solutions to mitigate oligotrophic conditions and increase biodiversity while maintaining water quality.

## MATERIALS AND METHODS

### Material

To monitor the cleanliness of Mikawa Bay and Ise Bay, water samples have been collected monthly from marine environmental reference points in Mikawa Bay and Ise Bay (Figure **S4a**) since April 1998, and the contents of transparency, salinity, DO (dissolved oxygen), COD (chemical oxygen demand), N (total nitrogen), P (total phosphorus), Chl-a, pheophytin, and each contaminant ion (Zn, Cd, Pb, Cr(VI), As, Hg, and anionic surfactants) have been measured. In particular, Mikawa Bay is managed in three separate marine areas, and standards for controlling total emissions for COD, N, and P are set for each area, as shown in Figure **S4a**. The simulated samples of the rivers were prepared using deionized water (pH: 5.0-6.0) as dilution water. Before preparing the samples, test tubes were sterilized by heating at 110 °C for 30 minutes. Each sample was placed in a test tube, tightly capped in air to mix the contents, and reacted at the prescribed temperature in the dark. All solvents and chemicals were of reagent grade quality, purchased commercially and used without further purification unless otherwise noted. As sodium chlorite, a chlorite ion standard solution for ion chromatography (ClO_2_^-^ ion concentration: 1000 ppm) manufactured by Fuji Film Wako Pure Chemical Industries, Ltd. was used.

### General

ESR spectra were measured with a Bruker-EMX Plus. The abundance of each ion in the simulated samples was measured using a Prominence nano equipped with anion-exchange and cation-exchange ion chromatograph, and each ion in the eluate was determined by conductivity or absorbance (at 210 nm). The ion exchanger used consisted of an anion separation column (Shim-pack IC-SA2) and a cation separation column (Shim-pack IC-C4), and the eluents were used Na_2_CO_3_/NaHCO_3_ aqueous solution and 18-crown-6 ether/CH_3_SO_3_H aqueous solution, respectively.

### ESR measurement

The hyperfine structure of the ESR spectra of radicals in sample water was measured. These samples were packed in a capillary tube and measured at room temperature. The ESR measurements were performed at a microwave frequency of 9.86 GHz, microwave power of 15.89 mW, and magnetic modulation amplitude of 0.5 GHz.

### Preparation of simulated samples

The samples were prepared by placing 1.0 mL of aqueous HNO_3_ (HNO_3_; 13.4 mmol/L, pH 2.0) and 1.0 mL of aqueous NaClO_2_ (14.8 mmol/L) in a test tube and sealing it in air. After mixing the contents of the sealed test tube, the samples were allowed to react at room temperature or 55 °C without stirring.

## RESULTS AND DISCUSSION

Ise Bay and Mikawa Bay, located adjacent to major urban and industrial centers, are increasingly affected by water pollution driven by anthropogenic activities. This has led to frequent occurrences of red tides and blue tides caused by eutrophication, severely impacting coastal fisheries. Efforts to mitigate environmental pollution, primarily through the purification of rivers and brackish lakes feeding into the bays, have yielded positive outcomes. As shown in Figure **S6a**, the number of red and blue tide occurrences has steadily decreased since 1990. However, since 1995, the number of red tide events has stagnated, despite continued nutrient decline. This stagnation can be attributed to nitrogen-fixing organisms such as cyanobacteria (12), which thrive by utilizing trace amounts of phosphorus, atmospheric nitrogen, and sunlight, making it challenging to completely eradicate red tides even in oligotrophic waters. The composition of phytoplankton communities in Mikawa Bay and Ise Bay is a dynamic system that changes rapidly depending on the season and environmental factors.(13) Diatoms generally dominate these waters, but as water temperatures rise and water mixing occurs, other microalgae—such as dinoflagellates and cyanobacteria— become more prominent, forming a diverse and complex community. In particular, dinoflagellates are typical species that cause red tides, which frequently occur along the coast of Japan, especially in Mikawa Bay and Ise Bay. While reductions in red tides have curtailed mass mortality events among marine organisms and decreased blue tide occurrences, some challenges persist. Although sand-covering has improved hypoxic conditions in brackish lakes (Figure **S6b**), its impact on further reducing blue tides remains limited. These findings suggest that a combination of seabed dredging and sand-covering is necessary for greater effectiveness. Additionally, decreasing pollution loads has transitioned the bays from eutrophic to oligotrophic conditions. Consequently, annual catches of benthic species, including mantis shrimp, conger eels, shrimp, and short-neck clams, have declined (Figures **S7a** and **S7b**). Similarly, catches of pufferfish, a higher-order consumer, have also decreased, despite the occurrence of a dominant year group (Figure S7c). These observations imply that a reduction in phytoplankton abundance disrupts the food chain, adversely affecting zooplankton and marine organism growth, ultimately impairing biodiversity. To address this, we focused on Chl-a concentration as an indicator of phytoplankton abundance, investigating strategies to improve trophic conditions while identifying the causes of oligotrophy in enclosed waters.

### Optimizing nutrient levels in enclosed seas: Setting target values for nutrient levels

In order to improve the nutrient condition of Mikawa Bay, we analyzed the impact of oligotrophy in enclosed waters on the fisheries industry. In the bay, the number of benthic organisms that feed mainly on phytoplankton, such as mantis shrimp, conger eels, and shrimp, has decreased since around 2011, and the number of tiger pufferfish, which rarely migrate outside the bay, has also decreased significantly since the same period. On the other hand, the catch of short-neck clams began to decrease around 1991, prior to the decrease in these benthic organisms and tiger pufferfish, and despite efforts such as transplanting and releasing juvenile clams, the catch of clams has continued to decrease significantly. Short-neck clams are considered a key indicator of marine oligotrophy, as their growth correlates closely with nutrient levels affecting phytoplankton. The degree of oligotrophy in Mikawa Bay was assessed using Chl-a concentration, a marker of phytoplankton abundance. To examine the impact of reductions in P and N on clam populations, we analyzed the relationship between clam density per unit area and Chl-a levels. Our analysis revealed that clam growth reached saturation at Chl-a concentrations of approximately 20 µg/L, while clam populations approached zero when Chl-a levels fell below 10 µg/L (Figure S8a). Furthermore, Figure **S8b** indicates that the annual average Chl-a concentration at the environmental reference point in Mikawa Bay has declined from 20 µg/L to below 10 µg/L. In addition, data from Tokyo Bay suggests that red tides may occur frequently when Chl-a concentrations exceed 50 µg/L, so it is desirable to set the target value of Chl-a concentration in Mikawa Bay between 20 µg/L and 50 µg/L.

### Enhancing Phytoplankton Growth in Mikawa Bay

P and N are essential macronutrients that support the biochemical processes and growth of phytoplankton in aquatic ecosystems. While it was initially hypothesized that sufficient levels of P and N in Mikawa Bay would promote phytoplankton growth, recent studies suggest that the optimal balance of N and P varies across trophic levels.(10,11) Literature data(14) indicate a weak correlation (Correlation Coefficient (CORR)=0.59) between declining N levels and reduced Chl-a concentrations (Figure **S9a**). In contrast, a strong positive correlation (CORR=0.93) exists between P and Chl-a, with reductions in P causing significant declines in Chl-a (Figure **S9b**). This aligns with earlier research on nutrient limitations in similar ecosystems(15), which identified phosphorus as a key factor driving phytoplankton biomass under eutrophic conditions. These findings suggest that the marine oligotrophy observed in Mikawa Bay is primarily attributed to reduced P levels rather than N, adversely affecting Chl-a concentrations.(16) To mitigate this issue, an intervention was implemented from 2017 to 2021 to increase P concentrations in treated water discharged from WWTPs into the bay. During this period, P levels in effluent from two WWTPs near Mikawa Bay were elevated from the usual 0.3 mg/L to approximately 0.7 mg/L for several months each year (Figures **S9c** and **S9d**). Despite these efforts, the enhanced P levels alone failed to yield significant improvements in clam catch or body size.(17)

### Reevaluating Nutrient Ratios for Phytoplankton Growth in Mikawa Bay: A Case Study on P and Chl-a Dynamics

To better understand these limited results, we reevaluated the relationship between P and Chl-a concentrations. Notably, even though P concentrations in WWTP effluent doubled throughout an 18-month period (6 months per year), the positive effects were confined to areas near the WWTP outfalls. This outcome underscores the ecosystem-wide limitations caused by unbalanced nutrient ratios. Prior studies(18) have shown that single-nutrient interventions often result in localized benefits but fail to address broader constraints. Furthermore, nonlinear relationships between nutrients and Chl-a were observed under oligotrophic conditions, with the P-Chl-a dynamics accurately described by a power regression model (Figure **S10a**). These findings highlight the necessity of a multifactorial approach to address nutrient imbalances and support phytoplankton growth effectively in Mikawa Bay. Specifically, when Chl-a was in the range of 15.38–303.20, the relationship between P and Chl-a was better fitted by power regression (coefficient of determination (R^2^) = 0.875, mean absolute error (MAE) = 0.010, P-value (P) = 0.258) than by linear regression (R^2^ = 0.858, MAE = 0.011, P = 0.143). Furthermore, the relationship between N and Chl-a was positively correlated when Chl-a was above 50 µg/L. However, fitting a linear model to the data showed that this correlation was almost negligible (R^2^ = 0.355, MAE = 0.073, P = 0.401). On the other hand, when Chl-a was less than 50 µg/L, a negative correlation was observed. This negative correlation was better explained by fitting a power regression curve to the data, which gave a stronger fit (R^2^= 0.777, MAE = 0.038, P = 0.499) (Figure **S10b**). These findings align with established models of nutrient dynamics (19), which emphasize non-linear relationships under nutrient-scarce conditions, reflecting the intricate dependencies within aquatic ecosystems. Our results on phytoplankton growth and nutrient ratios indicate that under oligotrophic conditions, phytoplankton growth depends on large amounts of N and relatively small amounts of P, and therefore increasing P alone has limited effect. Recent studies suggest that P supports phytoplankton growth under eutrophic conditions, whereas N serves as the primary limiting factor under oligotrophic conditions. (11) To explore this further, we conducted a multifactorial analysis examining the relationship between Chl-a, an established indicator of phytoplankton, and N and P. Our findings revealed a strong negative correlation between the N/P ratio derived from Chl-a concentrations and optimal N and P values (**Figure 1a**). Additionally, power regression (R^2^ = 0.953, MAE = 1.588, P = 0.318) demonstrated a superior fit compared to linear regression. Building on these results, we performed multiple regression analysis and response surface modeling, treating N and P as independent variables and Chl-a as the dependent response variable. This approach produced a three-dimensional response surface, shown in **Figure 1b**, which visually depicts the Chl-a levels corresponding to various N and P concentrations, and its top- and bottom-view projections in **Figures 1c** and **1d**. Lastly, the water quality data from Mikawa Bay and Ise Bay were used to validate the applicability of the relationships derived between P, N, and Chl-a.

**Figure 1.**
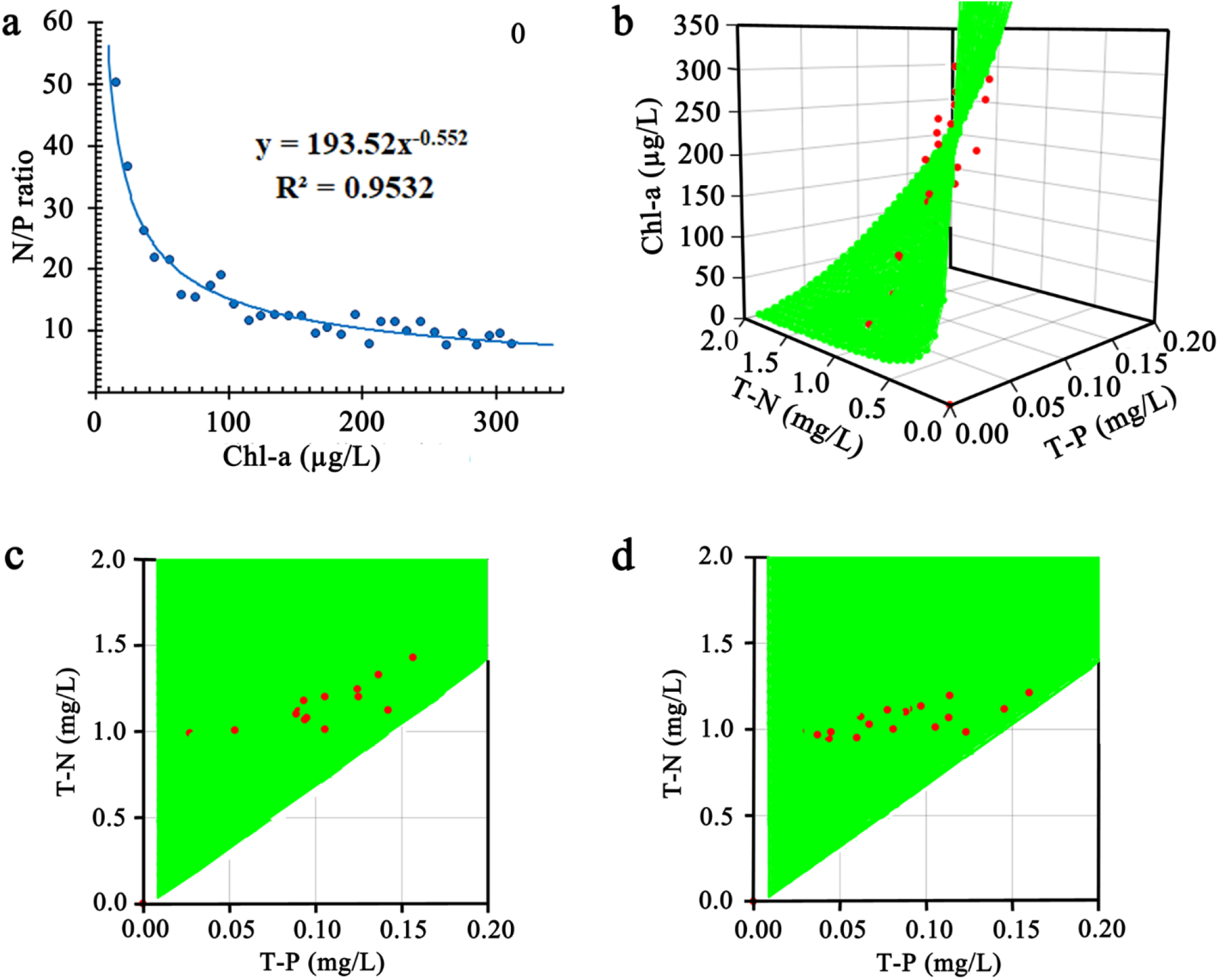
**a)** Correlation between N/P ratio and Chl-a and its response curve. **b**) Response surfaces were modelled on 3D scatter plots of Chl-a observations (red) using multiple regression analysis with P and N as explanatory variables and Chl-a as the response variable. **c**) Its top projection. **d**) Its bottom projection.

### Optimizing Nutrient Levels for Phytoplankton Growth: Insights from Multifactor Analysis in Ise and Mikawa Bays

Figure S11 illustrates the monthly relationships between P, N, or the N/P ratio, and Chl-a at nutrient-rich environmental reference points A-3, K-1, N-1, and K-2 in Ise Bay and Mikawa Bay from 1998 to 2023. At point A-3, unregulated agricultural wastewater inflow has resulted in significantly higher N concentration and overall better nutrient status compared to other points. In contrast, in Nagoya Port (N-1), which receives treated sewage water from Nagoya City, and Kinuura Port (K-1, K-2), which receives agricultural wastewater and septic tank effluent, the concentrations of P and N were lower than in A-3, and the decrease in N concentration was particularly significant. Despite these differences, Chl-a concentrations are almost the same across these points, suggesting that increased P concentration in eutrophic waters is the main driver of phytoplankton growth. Conversely, nutrient deficiency is becoming more pronounced at environmental reference points N-4, N-5, N-6, and N-7 in Ise Bay and Mikawa Bay. Figure **S12** shows that N concentrations at these points are lower than those estimated from their Chl-a values. The highest Chl-a values occur when both N and P concentrations align with the respective N vs. Chl-a and P vs. Chl-a response curves, indicating a position on the N/P ratio vs. Chl-a response curve. Under oligotrophic conditions, an increase in N concentration leads to a slight increase in Chl-a concentration, which subsequently increases the N/P ratio. Therefore, nitrogen availability appears to be the main limiting factor for phytoplankton growth in these areas. A social experiment is being conducted in Mikawa Bay, which is smaller than Ise Bay, to improve nutrient conditions. We compared monthly Chl-a measurements from 1998 to 2023 at four environmental reference points shown in Figure **S13** with Chl-a estimates obtained by substituting measured values of P and N into the response function from **Figure 1a**. **Figures 2a** and **2b** demonstrate that A-2, A-5, and A-13 had significantly lower P and N compared to A-3, yet there was no significant difference in Chl-a levels. These results suggest that Chl-a levels are highest when the N/P ratio aligns with the response curve. Conversely, Chl-a levels remain low even with high N and P levels when the N/P ratio deviates from this curve. The Chl-a concentration in Mikawa Bay indicates that N deficiency is the main limiting factor for phytoplankton growth. This phenomenon is also observed at other observation sites in Mikawa Bay and Ise Bay.

**Figure 2.**
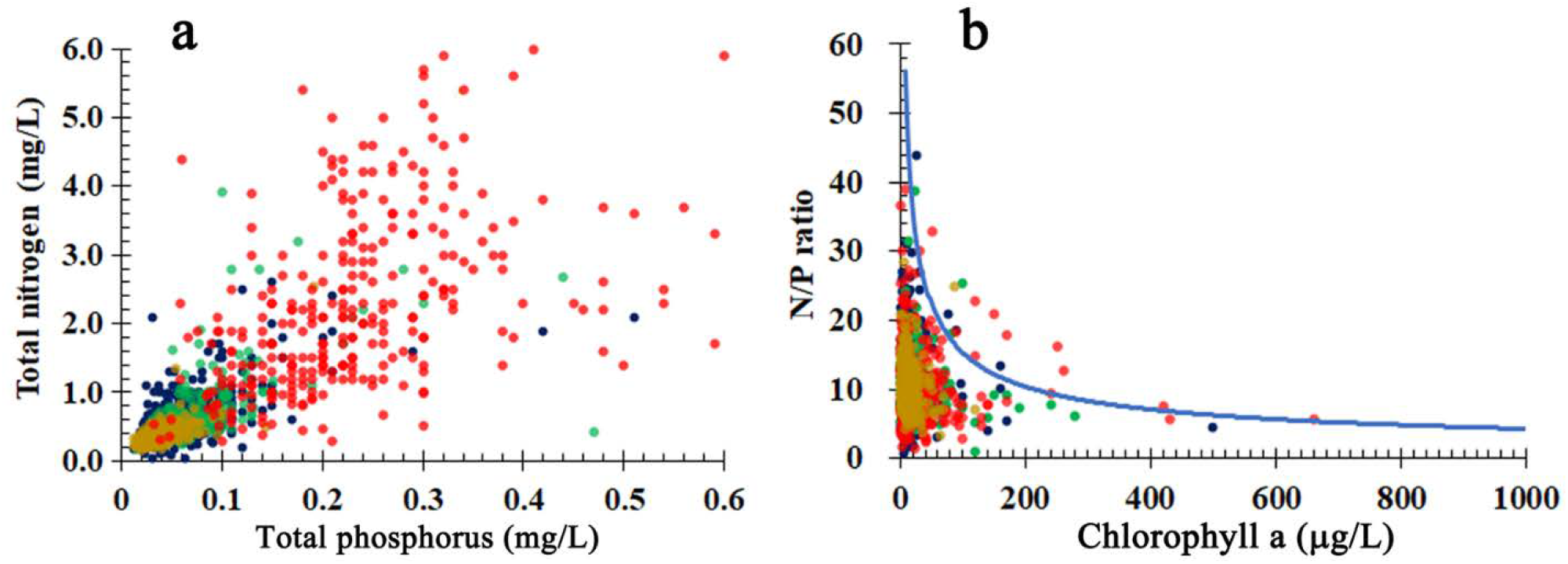
a) Scatter plots of observed values of samples taken at A-2, A-3, A-5, and A-13 every month from 1998 to 2023. P plotted on the horizontal axis and N on the vertical axis. b) Scatter plots of observed values plotted as Chl-a on the horizontal axis and N/P ratio on the vertical axis, and their response curves. Observed values at the environmental reference points are represented by A-2 (green), A-3 (red), A-5 (yellow), and A-13 (blue).

### Chemodenitrification: A Key Player in Nitrogen Cycling and Its Environmental Impact

This study investigates the N/P imbalance in Aichi Prefecture’s aquatic environments, focusing on the denitrification reactions that lead to nitrogen depletion. Using Electron Spin Resonance (ESR) measurements, ·NO and ·N radicals are identified as key intermediates. The research also explores electrochemical nitrate ion reduction to better understand N and P transformations. By integrating these approaches, the mechanisms behind oligotrophic conditions in marine ecosystems are revealed. In soils, most ammoniacal nitrogen in fertilizers is oxidized to NO_2_^-^ ions under aerobic conditions and subsequently to NO_3_^-^ ions, making NO_3_^-^ the predominant form of inorganic nitrogen. In soil water, dissimilatory nitrate reduction to ammonium (DNRA) occurs, where microorganisms oxidize organic matter and reduce NO_3_^-^ ions to NO_2_^-^ ions, then to NH_4_^+^ ions.(20,21) Assimilatory nitrate reduction also occurs, using NADH and reduced ferredoxin as electron donors, rather than NO_3_^-^ ions.(22,23) Recently, abiotic nitrate reduction

(chemodenitrification) by minerals and organic matter in soil water has also been reported, with Fe^2+^ ions in particular attracting attention.(24-29) This chemodenitrification leads to a decrease in inorganic nitrogen when it occurs in the ocean or in rivers that flow into the ocean. Figure **S14** shows the abundance of soluble Fe^2+^ and Fe^3+^ ions in surface waters receiving agricultural wastewater. Other metals also reduce nitrate, but none have been reported to oxidize NH_4_^+^ ions.(30-34) Our study showed that disinfection by-product ClO_2_^-^ ions in aqueous solutions act as chemical denitrification catalysts like Fe^2+^ ions in soil water, though they are less effective at oxidizing NH_4_^+^ ions.(35) Cyclic voltammetry studies revealed that the redox reaction between ·ClO_2_ radicals and ClO_2_^-^ ions is reversible, with redox potentials reported at E^0^ = 0.700 V vs SCE(36) and recently updated to E^0^ = 0.954 V vs SCE.(37) With Fe^3+^ ions/Fe^2+^ ions and NO_3_ ^-^ ions/NO_2_^-^ ions redox potentials at 0.77 V and 0.835 V respectively, Fe^2+^ ions are unlikely to reduce NO_3_^-^ ions electrochemically. Conversely, the redox potential of ·ClO_2_ radical/ClO_2_^-^ ion suggests that ClO_2_^-^ ion can catalyze nitrate reduction. Furthermore, by focusing on the generation process of ·NO radical and ·N radical, which are thought to be intermediates in the denitrification reaction, it is expected that the gap between wastewater treatment processes and the nitrogen cycle in the environment will be clarified, leading to the development of a new electrochemical approach. **Figure 3a** demonstrates that the reduction of NO_3_^-^ ions is driven by the oxidation of ClO_2_^-^ ions to ·ClO_2_ radicals. Although chemical denitrification has not been reported in aerobic rivers or marine surface waters, water quality data from activated sludge and chlorination treatments show oxidation of NH_4_^+^ ions and accumulation of NO_3_^-^ ions. This suggests that chemical denitrification may occur under certain artificial conditions and may affect aquatic environments receiving treated effluents. However, further investigations are needed to clarify its role in natural aquatic systems. **Figure 3b** highlights the oxidation of ClO_2_^-^ ions to ·ClO_2_ radicals, followed by a subsequent decline in ·ClO_2_ radicals as the reaction reached completion. **Figure 3c** shows that at room temperature, more than 90% of NO_3_^-^ ions were reduced, while over 60% of ClO_2_^-^ ions remained intact. Trace amounts of Cl^-^ and ClO_3_^-^ ions were also detected, pointing to the potential formation of chlorine nitrate (ClNO_3_), as supported by the scavenging abilities of ·ClO_2_ radicals for ·NO radicals.(38,39) No cations other than Na^+^ ions were observed (**Figure 3e**). At 55°C (**Figures 3d and 3f**), more than 90% of NO_3_^-^ ions were reduced, while over 60% of ClO_2_^-^ ions persisted. Slight increases in Cl^-^ and ClO_3_^-^ ions and traces of NH_4_^+^ ions were detected, suggesting partial reduction of NO_3_^-^ ions to NH_4_^+^ ions. The presence of ·NO and ·N radicals, confirmed using spin-trapping agents, indicates the release of nitrous oxide (N_2_O) and nitrogen gas (N_2_). As a result of these coupling reactions, only trace amounts of nitrogen components were detected by ion chromatography. These radicals are released into the atmosphere as N_2_O and N_2_ produced by coupling reactions. The inability to observe ·ClO_2_ radicals in ESR experiments involving spin trap agents (40,41) is likely attributable to the short lifetime of ·ClO_2_ radicals compared to PTIO (2-phenyl-4,4,5,5-tetramethylimidazoline-3-oxide-1-oxyl).(42) Unlike dissimilatory nitrate reduction to NH_4_^+^ ion and assimilatory nitrate reduction, abiotic chemical denitrification reduces inorganic nitrogen by producing N_2_O and N_2_ gases that are subsequently released into the atmosphere. In oligotrophic closed sea areas, this process lowers N nutrient levels without affecting P nutrients, thereby decreasing the N/P ratio.

**Figure 3.**
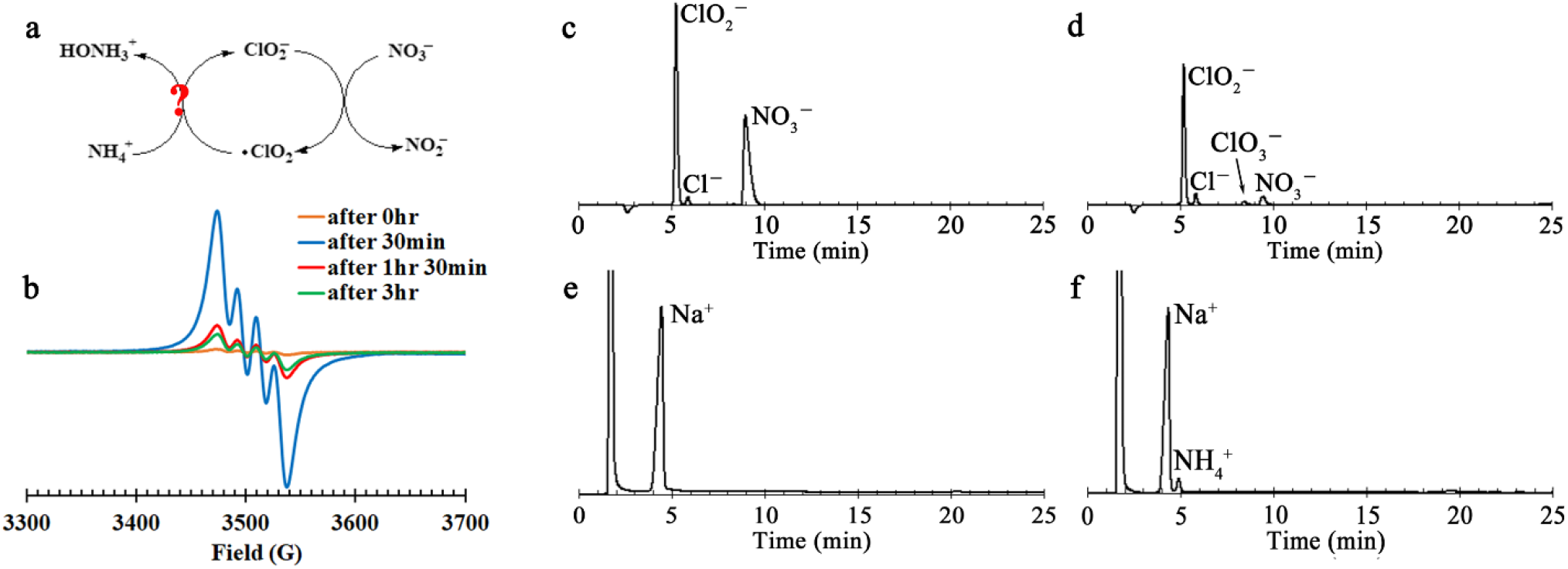
**a)** Schematic diagram of the effect of ClO_2_^-^ ions in promoting the decomposition reaction of NH_4_NO_3_. **b)** The progress of the reaction was monitored in an aqueous solution (pH 2.2) containing NaClO_2_ (7.41 mmol/L) and HNO_3_ (6.57 mmol/L) at 55 °C by ESR spectroscopy of the ClO_2_ radical reduced from the ClO_2_^-^ ion. **c)** Anion composition at time of sample preparation reacted at room temperature. **d)** Anion composition after 24 hours of reaction at room temperature. **e)** Cation composition after 24 hours of reaction at room temperature. **f)** Cation composition after 24 hours of reaction at 55°C.

### Balancing Nutrient Levels to Prevent Red Tides

An analysis of P and N concentration distribution data from the 1990s to the 2020s in Mikawa Bay and Ise Bay shows a consistent decline in these concentrations (Figures **S5a-S5d**). In Mikawa Bay, observed P levels exceed predictions based on Chl-a levels (Figures **S13d, S13g, S13j**), while observed N levels are lower than predicted (Figures **S13e, S13h, S13k**). Currently, Chl-a levels in both Mikawa Bay and Ise Bay reflect oligotrophic conditions caused by nitrogen deficiency, which limits phytoplankton growth. Even in A-3, where unregulated agricultural runoff has an impact, Chl-a concentrations remain low due to an imbalance in the N/P ratio, even though P and N concentrations are significantly high (Figures. **S13a**–**S13c**). Figure 4a illustrates that the measured Chl-a levels at the non-oligotrophic point A-3 generally remained within the boundaries of the three-dimensional response surface, which was modeled using N and P as independent variables. Table **S1** confirms that, across various environmental reference points, the Chl-a levels rarely exceeded the modeled response surface defined by the input of N and P values. Outlier rates (only observations falling on the upper side of the response surface) were consistently less than 0.04 for all environmental reference points and generally less than 0.015 when environmental anomalies such as red tides were excluded. Furthermore, the 95% confidence intervals for the population outlier rate did not exceed 0.06, and were less than 0.025 when environmental anomalies were excluded. The 3D scatter plots of points A-3 and A-13 (**Figures 4a** and **4b**), represented using the projection method (Figures **S15** and **S16**), demonstrate how Chl-a response surfaces shift under varying P and N concentrations. **Figures 4c** and **4d** further reveal that maintaining optimal Chl-a levels (20–50 µg/L) requires adjusting P and N concentrations to specific ranges. This multiple regression response surface analysis enables a detailed examination of fine-scale variations in Chl-a levels. Response surface 2 (**Figure 4d**) highlights that Chl-a levels remain within the optimal P and N ranges defined by the regression analysis of P vs. Chl-a and N vs. Chl-a. In contrast, response surface 1 (**Figure 4c**) displays a narrower range of variation. Response surface 1 indicates that the lowest P concentration combined with the highest N concentration results in the minimum Chl-a value (20 µg/L), while the highest P concentration paired with the lowest N concentration leads to the maximum Chl-a value (50 µg/L). The broader ranges shown in response surface 2 suggest a potential risk of Chl-a levels exceeding the desired range (20–50 µg/L) depending on the specific concentrations of P and N used. Based on the response surface analysis, the suitable concentration ranges for P and N were determined to be 0.026–0.046 mg/L and 0.95–1.08 mg/L, respectively. To mitigate the risk of harmful red and blue tides, it is recommended to maintain P and N concentrations within these ranges. This study builds upon earlier findings (43) by offering a refined understanding of the nutrient thresholds required to balance healthy phytoplankton growth while minimizing ecological risks.

**Figure 4.**
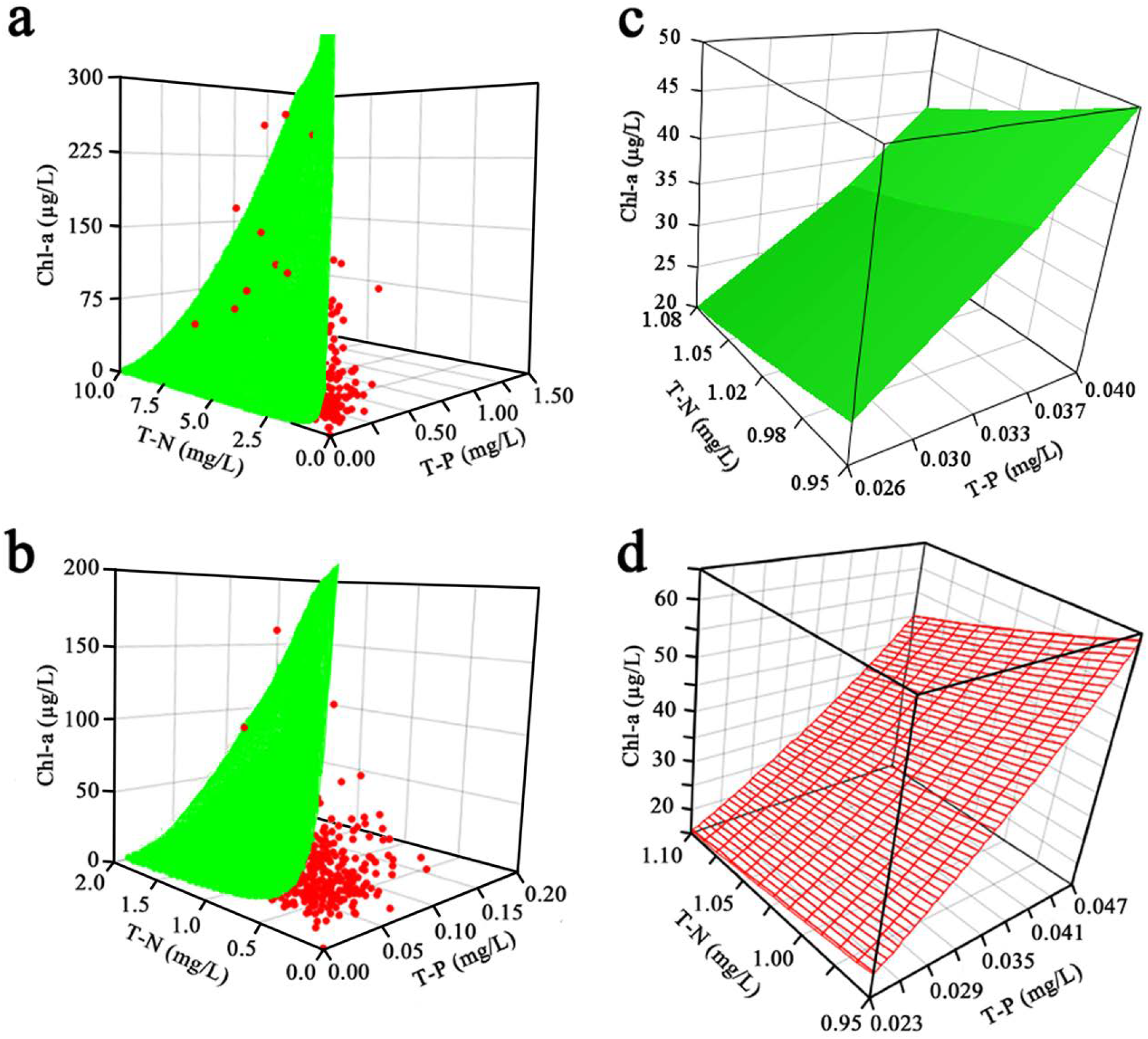
Observed values (red circles) of samples collected monthly from 1998 to 2023 at environmental reference points (A-3, A-13) and their response surface scatter plots are plotted in a three-dimensional coordinate system with P and N as explanatory variables and Chl-a as the objective variable. a) A-3. b) A-13. c) Response surface 1 for Chl-a values ranging from 50 µg/L to 20 µg/L. d) Response surface 2 for Chl-a values ranging from 50 µg/L to 20 µg/L obtained from the respective response curves of P-Chl-a and N-Chl-a shows that response surface 2 extends beyond the range of 50 µg/L to 20 µg/L.

### Nutrient supply strategy in enclosed sea areas

Currently, there is no established method to uniformly increase the amount of N throughout Mikawa Bay. For example, it has been shown that nutrients can be replenished at environmental reference points A-3 and N-1 by mixing neutral nitrogen fertilizer with agricultural and urban wastewater from land. In addition, it is possible to uniformly supply nutrients to each point in the bay by adjusting the discharge amount and nutrient concentration at existing WWTPs. Another method to immediately improve the oligotrophic state is to adjust the N concentration by adding appropriate amounts of nitrogen-containing fertilizer or organic matter to specific points, but this method requires careful operation as it may cause an imbalance in the N concentration. On the other hand, if our hypothesis is correct, even if the amount of nitrogen is excessive, it may be possible to suppress the occurrence of blue-green algae if the P concentration can be sufficiently reduced. Based on this, as a long-term measure, it is effective to suppress the nitrogen concentration by supplying nutrients from land areas and WWTPs. In addition, if the nitrogen concentration in the treated water from WWTPs discharged into Mikawa Bay increases, a N concentration gradient will occur near the discharge outlet, but since this effect is localized, the risk of red tide occurrence is thought to be further reduced (Figure **S17**).

## CONCLUTION

Ise Bay and Mikawa Bay are enclosed sea areas that frequently experience red and blue tides. Purification measures have been implemented to reduce COD, P, N, and Chl-a levels. However, these measures have led to oligotrophy in both bays. Sudden increases in P or N to address nutrient shortages could raise the risk of red and blue tides, so careful management is needed. To address nutrient shortages effectively, it is essential to identify the causes and minimize fluctuations in P and N levels. Our study focused on the correlation between Chl-a and the N/P ratio to find ways to safely increase Chl-a. We found that slightly reducing the current P levels could prevent Chl-a from exceeding 50 µg/L, even if N levels increased significantly. In the long term, increasing the nitrogen content in WWTP effluent by twofold could raise Chl-a levels to over 20 µg/L. The increase is expected to enhance habitats for benthic organisms and zooplankton, promote the re-establishment of phytoplankton-based food webs, and help maintain biodiversity in enclosed sea areas.

## Supporting information

Purification Efforts of Aburagafuchi-Lake to Combat Red and Blue Tides. (Figures S1c, S1e, S2a, and S2b); Nutrient Dynamics in Aburagafuchi: Trends an

## Author Contributions

I.N. conceived the study, designed the study, supervised study, and conducted the sampling. I.N. and S.I. performed the ESR measurements and analysis. I.N. and K.N. collected information on oligotrophy in the waters around Japan. The manuscript was written through contributions of all authors. All authors have given approval to the final version of the manuscript.

## Notes

The author declares no competing financial interest.

## ACKNOWLEDGMENT

This research result is part of the results of a survey conducted by Aichi Prefecture in Mikawa Bay and Ise Bay, and we would like to thank the relevant staff for their helpful advice. We thank Kaito Kiritoshi, Mituhiro Gonda, Chika Makino, Hideaki Makihara, Mamoru Yasui, Chikako Inoue, Minoru Oda, Takanori Matui, Yoshihiro Yokoi, Seigo Horie, Rie Okuda, Takeru Sintani, Hiroshi Watanaba, Takae Kawai (Bureau of the Environment, Aichi Pref.) for assistance in the field. We thank Dr. Tomoyuki Shikata (Fisheries Technology Institute) for providing information on oligotrophy in the Seto Inland Sea. We thank Professor Toshihiko Yokoyama (Institute for Molecular Science) for helpful discussions. We acknowledge the efforts of Dr. Mizue Asada and Motoki Fujiwara (Institute for Molecular Science) in ESR EMX Plus measurements, data collection, and analytical evaluation. We acknowledge the efforts of Haruyo Nagao (Institute for Molecular Science) in JNM-ECA600 measurements, data collection, and analytical evaluation. This work was conducted in Institute for Molecular Science, supported by “Advanced Research Infrastructure for Materials and Nanotechnology in Japan (ARIM)” of the Ministry of Education, Culture, Sports, Science and Technology (MEXT). Proposal Number JPMXP1223MS1097.

## ABBREVIATIONS

PTIO: 2-Phenyl-4,4,5,5-tetramethylimidazoline-3-oxide-1-oxyl ; 18-Crown 6-Ether, 1,4,7,10,13,16-Hexaoxacyclooctadecane

## SYNOPSIS TOC

The impact of oligotrophication of enclosed seas on the food chain within the bay.

## CONTENTS OF SUPPORT INFORMATION

Purification Efforts of Aburagafuchi-Lake to Combat Red and Blue Tides. (Figures **S1c, S1e, S2a**, and **S2b**); Nutrient Dynamics in Aburagafuchi: Trends and Implications. (Figure **S3**); Environmental Reference Points in Ise Bay and Mikawa Bay, and Annual Changes in Water Quality in Mikawa Bay. (Figure **S4**); Comparative Analysis of Nutrient and Phytoplankton Concentration Distributions in Mikawa Bay and Ise Bay Over Four Decades. (Figure **S5**); Temporal Trends in Red and Blue Tide Events in Ise Bay and Mikawa Bay and Mitigation Strategies for Blue Tides in Aburagafuchi-lake. (Figure **S6**); Annual Changes in Catch Volume in Ise Bay and Mikawa Bay (mainly Mikawa Bay). (Figure **S7**); Impact of Chlorophyll a Concentration on Clam Growth in Mikawa Bay. (Figure **S8**); Details of the Social Experiment Conducted in Mikawa Bay by the Aichi Prefectural Government. (Figure **S9**); Rethinking Nutrient Dynamics and Chlorophyll a Correlation Under Oligotrophic Conditions. (Figure **S10**); Phosphorus as the Key Driver for Phytoplankton Growth in Eutrophic Waters of Ise Bay and Mikawa Bay: A 25-Year Study. (Figure **S11**); Nitrogen Limitation as the Primary Factor for Phytoplankton Growth in Nutrient-Deficient Areas of Ise Bay and Mikawa Bay: A 25-Year Study. (Figure **S12**); Unexpected Chl-a Levels in Mikawa Bay: Discrepancies Between Nutrient Concentrations and Phytoplankton Growth. (Figure **S13**); Iron Ions in the Surface Water of the Rivers Flowing into Mikawa Bay may be a Potential Source of Metal Ions for Chemodenitrification. (Figure **S14**); Anomalies in Chl-a Concentration and Assessment of Environmental Equilibrium in Ise Bay and Mikawa Bay. (**Table S1**); Developed Views of the 3D Scatter Plots of A-3 and A13 in **Figures 4a** and **4d**. (Figures **S15** and **S16**); Investigating Nutrient Concentration Changes near the Wastewater Treatment Plant (WWTP) Outlet. (Figure **S17**)

