## Supplementary material for "From Red Tides to Healthy Ecosystems by Nutrient Management in Ise Bay and Mikawa Bay": Purification Efforts of Aburagafuchi-Lake to Combat Red and Blue Tides. (Figures S1c, S1e, S2a, and S2b); Nutrient Dynamics in Aburagafuchi: Trends an

### Support Information

### Contents

1. Purification Efforts of Aburagafuchi-Lake to Combat Red and Blue Tides. (Figures **S1c**, **S1e**, **S2a**, and **S2b**) ・・・・・pages2~4
2. Nutrient Dynamics in Aburagafuchi: Trends and Implications. (Figure **S3**) ・・・・・pages4~5
3. Environmental Reference Points in Ise Bay and Mikawa Bay, and Annual Changes in Water Quality in Mikawa Bay. (Figure **S4**) ・・・・・page 5
4. Comparative Analysis of Nutrient and Phytoplankton Concentration Distributions in Mikawa Bay and Ise Bay Over Four Decades. (Figure **S5**) ・・・・・page 6
5. Temporal Trends in Red and Blue Tide Events in Ise Bay and Mikawa Bay and Mitigation Strategies for Blue Tides in Aburagafuchi-lake. (Figure **S6**) ・・・・・page 7
6. Annual Changes in Catch Volume in Ise Bay and Mikawa Bay (mainly Mikawa Bay). (Figure **S7**) ・・・・・page 7
7. Impact of Chlorophyll a Concentration on Clam Growth in Mikawa Bay. (Figure **S8**) ・・・・・pages 8~9
8. Details of the Social Experiment Conducted in Mikawa Bay by the Aichi Prefectural Government. (Figure **S9**) ・・・・・page 9
9. Rethinking Nutrient Dynamics and Chlorophyll a Correlation Under Oligotrophic Conditions. (Figure **S10**) ・・・・・page 10
10. Phosphorus as the Key Driver for Phytoplankton Growth in Eutrophic Waters of Ise Bay and Mikawa Bay: A 25-Year Study. (Figure **S11**) ・・・・・pages 10~11
11. Nitrogen Limitation as the Primary Factor for Phytoplankton Growth in Nutrient-Deficient Areas of Ise Bay and Mikawa Bay: A 25-Year Study. (Figure **S12**) ・・・・・pages 11~12
12. Unexpected Chl-a Levels in Mikawa Bay: Discrepancies Between Nutrient Concentrations and Phytoplankton Growth. (Figure **S13**) ・・・・・pages 12~13
13. Iron Ions in the Surface Water of the Rivers Flowing into Mikawa Bay may be a Potential Source of Metal Ions for Chemodenitrification. (Figure **S14**) ・・・・・page 14
14. Anomalies in Chl-a Concentration and Assessment of Environmental Equilibrium in Ise Bay and Mikawa Bay. (**Table S1**) ・・・・・page 15
15. Developed Views of the 3D Scatter Plots of A-3 and A13 in Figures **4a** and **4d.** (Figures **S15** and **S16**) ・・・・・pages 15~17
16. Investigating Nutrient Concentration Changes near the Wastewater Treatment Plant (WWTP) Outlet. (Figure **S17**) ・・・・・pages 17~18

#### 1. Purification Efforts of Aburagafuchi-Lake to Combat Red and Blue Tides. (Figures **S1c**, **S1e**, **S2a**, and **S2b**)

Purification of Aburagafuchi-lake, a brackish lake connected to Mikawa Bay, is effective in preventing red and blue tides in enclosed water areas such as Mikawa Bay and Ise Bay. Figures **S1a** and **S1b** show that Aburagafuchi-lake, once the second most polluted brackish lake in Japan, is adjacent to automobile and agricultural zones. Since 1992, Aichi Prefecture and neighboring municipalities have been working to improve its water quality to meet national environmental standards.

Initially, efforts focused on preventing pollutant discharge at the source. Figures **S1d** and **S1c** indicate that the population around the lake is growing due to the profitable automobile industry, with domestic wastewater

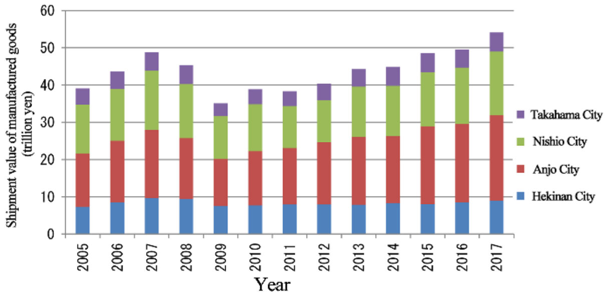


a, Changes in shipment value of manufactured in 4 cities in the Aburagafuchi basin.

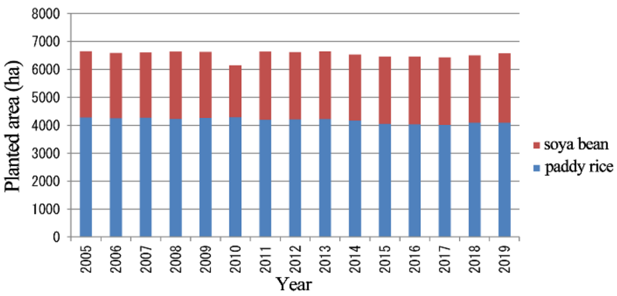


b, Changes in major planted areas in 4 cities in the Aburagafuchi basin.

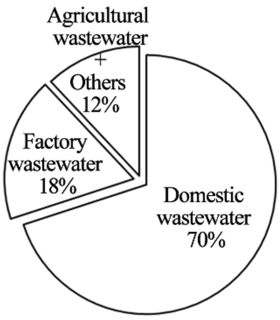


c, Percentage of pollutant load in Abragafuchi (2018).

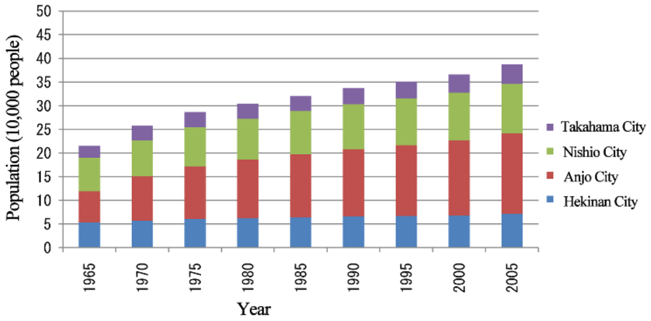


d, Changes in the population of the 4 cities around the Aburagafuchi.

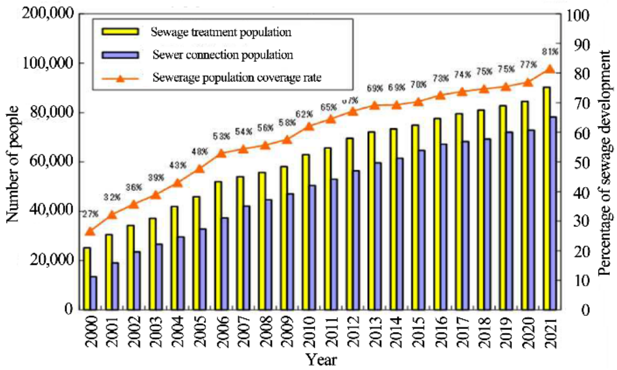


e, Changes in population and number of households connected to the sewage system around Aburagafuchi

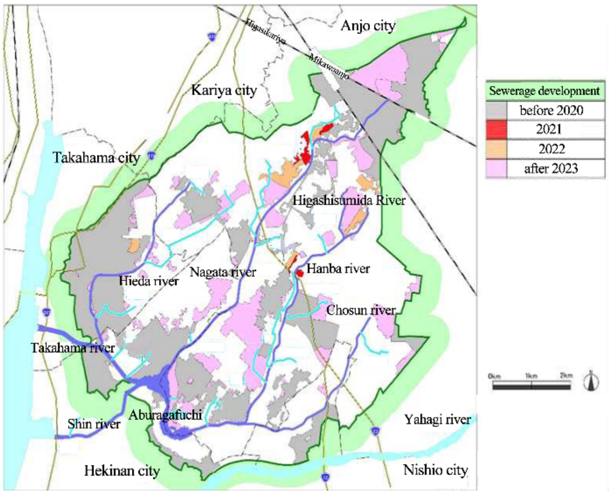


f, Progress of Sewerage Development around Aburagafuchi

**Figure S1.** Changes in the urban environment of the 4 cities around Aburagafuchi.

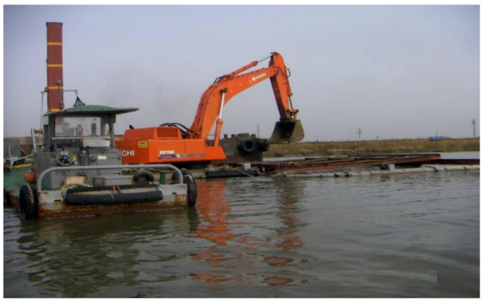

a, Dredging and sand-covering work in Abragafuchi.

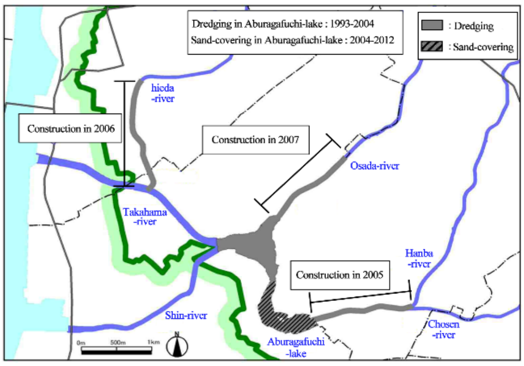


b, Construction periods and areas of dredging and sand-covering in Aburagafuchi.

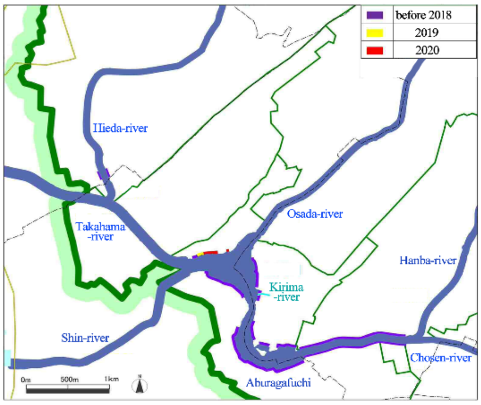


c, Areas and periods of vegetation improvement on the lakeshore of Aburagafuchi.

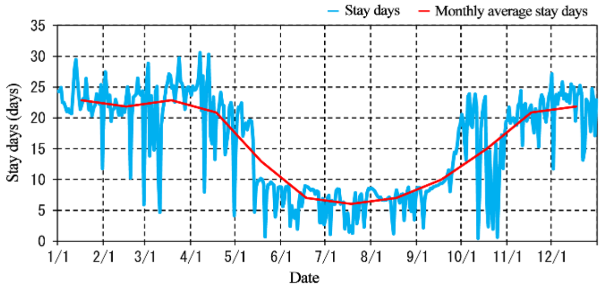


d, Dwell time of water in Aburagafuchi-lake estimated from flow rate of inflowing river (2019).

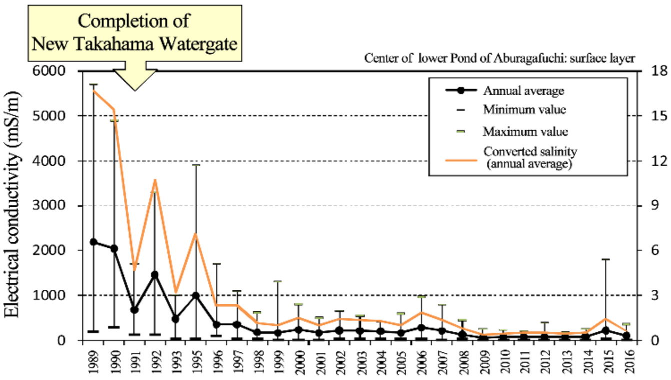


e, Annual mean salinity estimated from electrical conductivity of surface water in Aburagafuchi.

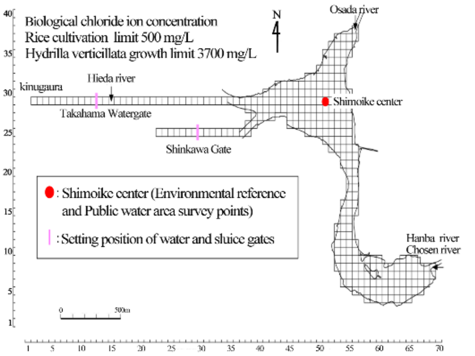


We divided Aburagafuchi into 50m horizontal grids and 6 vertical layers (1st layer: 0~2m, 2nd layer: 2~2.5m, 3rd layer: 3~3.5m, 4th layer: 3~3.5m, 5th layer: 3.5~4m, 6th layer: 4m~), and calculated the flow velocity, water temperature, and salinity of each grid.

f, Simulation salinity at subdivided points in Aburagafuchi.

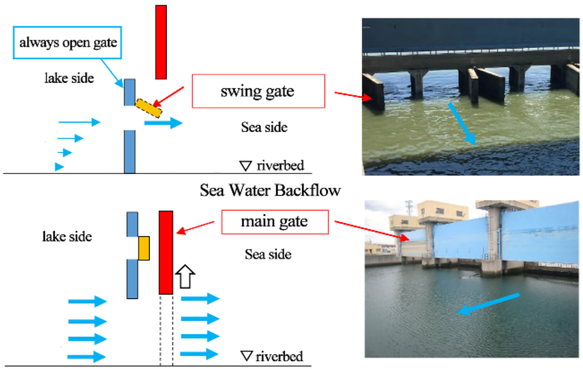


g, By opening and closing the main gate of the Takahama Floodgate, the old Aburagafuchi lake water is drained and new water is introduced into the river.

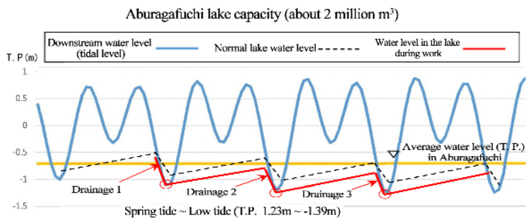


Opening closing operation of the Tkahama watergate
- Main gate opens at the average waterin the lake
- Drainage at low tide
- Main gate closed at low tide

h, Changes in the amount of lake water in Aburagafuchi by opening and closing the main gate of Takahama watergate

**Figure S2.** Progress of Water Purification of Aburagafuchi.

contributing about 70% of the lake's pollution load. However, Figures **S1e** and **S1f** demonstrate that by 2019, sewerage coverage reached 76%, leading to a steady decline in domestic wastewater pollution, with ongoing efforts to reach 100% coverage. Despite strict industrial wastewater regulations, agricultural wastewater pollution remains high due to insufficient legal restrictions. Overall, the expansion of sewage systems has significantly reduced nutrient inflow into enclosed water bodies.

Subsequently, dredging and sand-capping were conducted to remove pollutants from microorganisms deposited in the lake's hyporheic zone and river mouths. Before dredging, the sediment mainly comprised over 90% silt and clay with sulfides, indicating organic contamination. Post sand-capping, the proportion of sulfide-containing silt and clay decreased. Figures **S2a** and **S2b** show that approximately 110,000 m² of dredging and 40,000 m² of sand-capping were executed. Reed regeneration and embankment construction aimed at a more natural lakeshore, with Figure **S2c** showing 70% greening of the total lakeshore area.

Figure **S2d** shows that the Aburagafuchi-lake is a highly closed brackish water lake, so water stays in the lake for a long time. Figure **S2e** reveals that the lake's salinity is controlled by the Takahama Watergate. Consequently, as Figure **S2f** simulations suggest, attempts were made to replace old lake water with new river water while maintaining the lake's salinity. Figures **S2g** and **S2h** illustrate that the Takahama floodgate can open to drain old lake water to the sea and introduce new river water. **Table S1** confirms that this operation is regularly performed to ensure continuous water renewal in Aburagafuchi-lake.

**Table S1.** The amount of water discharged into the sea of Aburagafuchi-lake by opening and closing the main gate of Takahama Floodgate (x 10000m$^3$). Aburagafuchi-lake capacity (about 200$\times$10000m$^3$).

| Month | | | 4 | 5 | 6 | 7 | 8 | 9 | 10 | 11 | 12 | 1 | 2 | 3 | sum |
|---|---|---|---|---|---|---|---|---|---|---|---|---|---|---|---|
| 2020 | Discharge | $\times$10000m$^3$ | 75 | 81 | 295 | 511 | 347 | 321 | 71 | 132 | 70 | 144 | 99 | 124 | 2270 |
| | Exchange rate | % | 38 | 41 | 148 | 256 | 174 | 161 | 36 | 66 | 35 | 72 | 50 | 62 | |
| 2021 | Discharge | $\times$10000m$^3$ | 202 | 328 | 310 | 342 | 430 | 320 | 178 | 46 | 117 | 98 | 75 | 143 | 2619 |
| | Exchange rate | % | 101 | 164 | 155 | 171 | 215 | 160 | 89 | 38 | 59 | 49 | 38 | 72 | |

#### 2. Nutrient Dynamics in Aburagafuchi: Trends and Implications. (Figure S3)

Figure **S3a** shows that the nitrogen (N) content of the surface water in Aburagafuchi, the only brackish lake in Aichi Prefecture, has significantly decreased since 1998, but has remained almost constant since 2011. This stagnation is likely due to a slowdown in the rate of sewerage coverage increase, which now exceeds 65%. On the other hand, Figure **S3b** indicates that phosphorus (P) levels have hardly decreased since 1998. This phenomenon

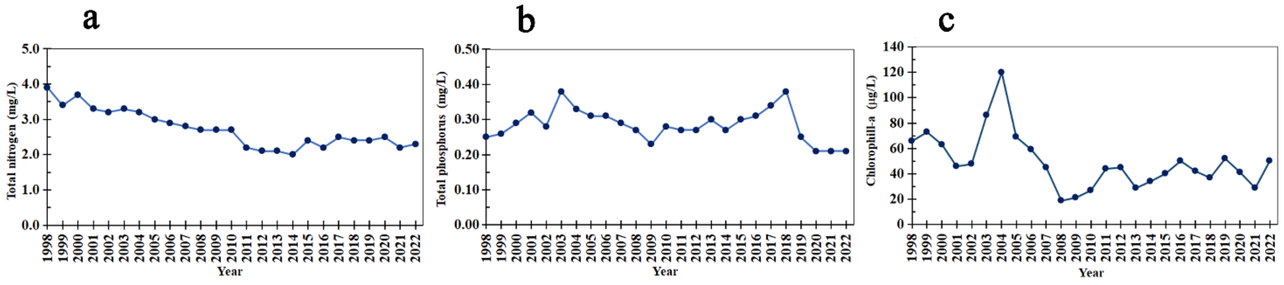


**Figure S3.** Annual changes in water quality over time in the Aburagafuchi surface water. **a)** Changes in nitrogen content over time. **b)** Changes in phosphorus content over time. **c)** Changes in chlorophyll a content over time.

is likely because there are no wastewater treatment plants (WWTPs) around Aburagafuchi, and only treated water from individual and collective septic tanks flows into the lake. However, Figure **S3c** shows that the reduction in nitrogenous nutrients due to this purification process has led to a decrease in chlorophyll-a (Chl-a), an indicator of phytoplankton, in Aburagafuchi.

#### 3. Environmental Reference Points in Ise Bay and Mikawa Bay and Annual Changes in Water Quality in Mikawa Bay. (Figure S4)

Long-term changes in water quality are being investigated in the enclosed waters of Ise Bay and Mikawa Bay, as shown in Figure **S4a**. Mikawa Bay, known for having the highest clam catch in Japan, is divided into three monitoring areas to observe changes in nitrogen (N) and phosphorus (P).

Figures **S4b** and **S4c** illustrate that N and P levels in Mikawa Bay (II) and Mikawa Bay (III), both rich in clams, exhibit minimal interannual fluctuation. This stability can be attributed to the dilution effect of seawater, which masks variations in N, P, and chlorophyll-a (Chl-a) at many observation points in these central parts of the bay. In contrast, Mikawa Bay (IV), being smaller in size, shows an annual reduction in N and P due to purification efforts in Aburagafuchi and surrounding areas. Furthermore, Figure **S4d** shows that Chl-a has decreased significantly.

Additionally, agricultural wastewater from points A-3 and A-12, which is not subject to legal regulation, flows into Mikawa Bay (II), resulting in higher P and N concentrations compared to Mikawa Bay (III). Despite this, the Chl-a levels in both areas were almost the same. The P and N levels in Mikawa Bay (IV) decreased to levels similar to those in Mikawa Bay (II) but did not reach the lower levels found in Mikawa Bay (III), likely due to the limitations of the current sewage treatment capacity.

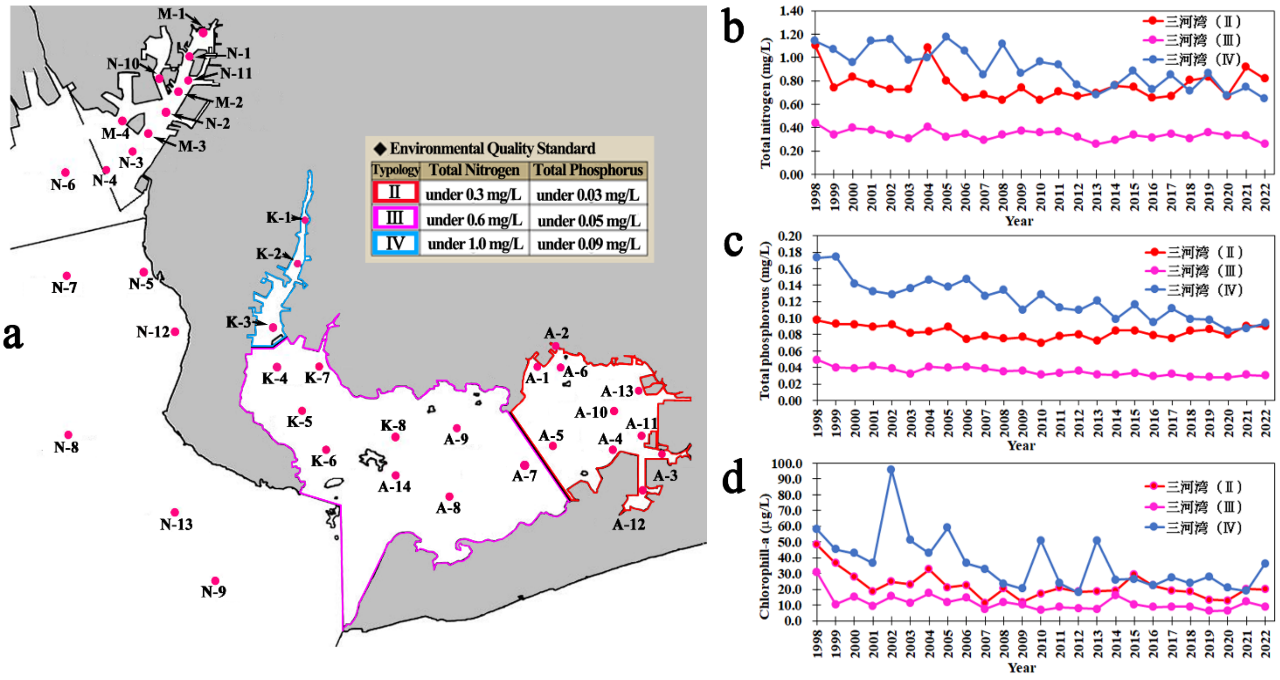


Figure **S4**. **a**) The marine environmental reference points of Mikawa Bay and Ise Bay, and the marine areas of Mikawa Bay divided into three environmental management zones. **b**) Changes in N concentration over time in each area of Mikawa Bay. **c**) Changes in P concentration over time in each area of Mikawa Bay. **d**) Changes in Chl-a concentration over time in each area of Mikawa Bay.

#### 4. Comparative Analysis of Nutrient and Phytoplankton Concentration Distributions in Mikawa Bay and Ise Bay Over Four Decades. (Figure S5)

Annual mean values of P, N, and Chl-a detected at various environmental reference points in Mikawa Bay (Figures **S4b**, **S4c**, and **S4d**) did not clearly show overall changes in water quality. Thus, we compared the average concentration distributions of N, P, and Chl-a in Mikawa Bay and Ise Bay from the periods 1980-1989 and 2010-2019. The results revealed that N concentrations were diluted from the 1980s (Figure **S5a**) to the 2010s (Figure S**5b**), P concentrations similarly decreased from the 1980s (Figure **S5c**) to the 2010s (Figure **S5d**), and Chl-a concentrations, indicating phytoplankton levels, also dropped from the 1980s (Figure **S5e**) to the 2010s (Figure **S5f**).

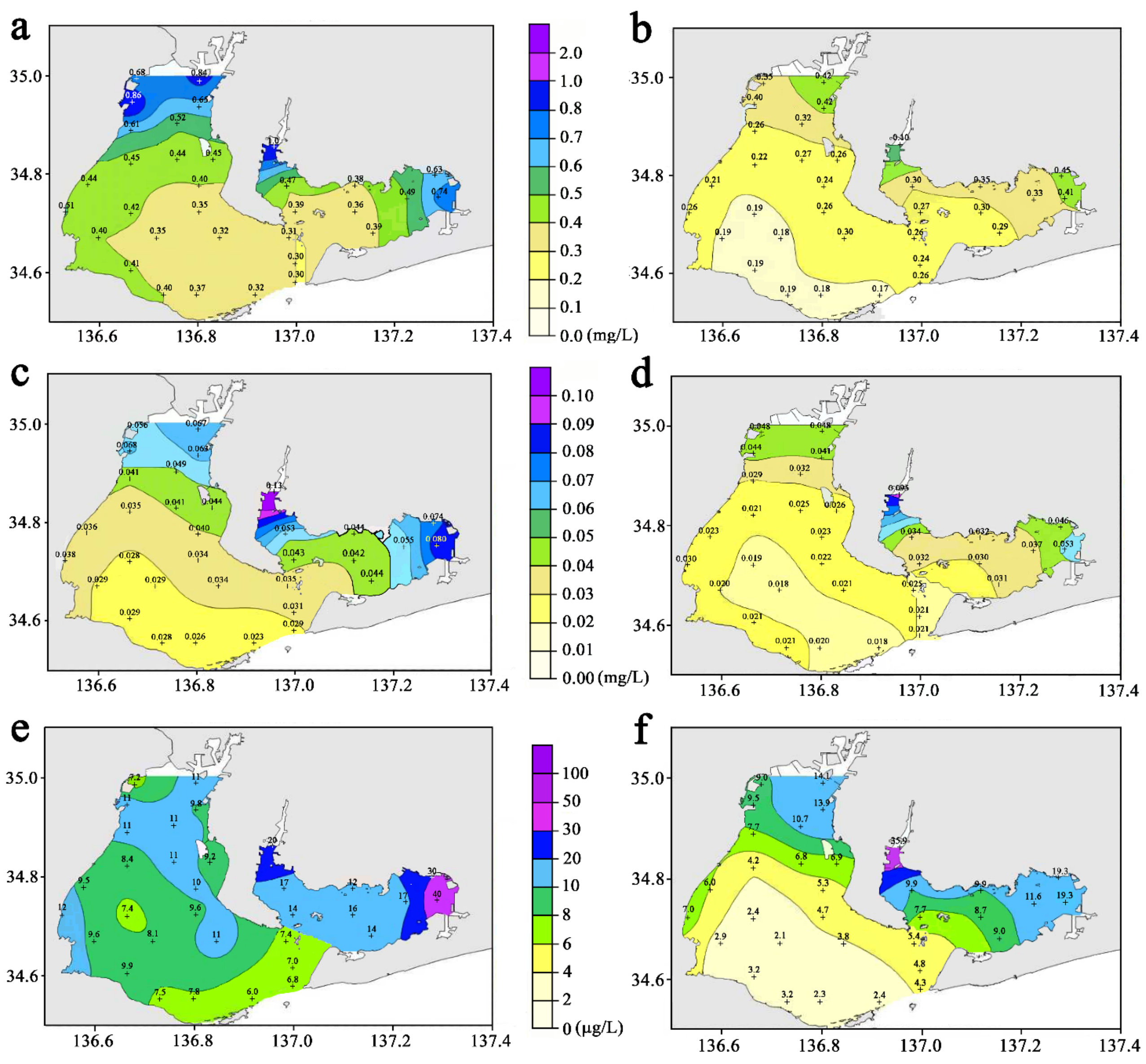


Figure **S5.** Comparison of annual water quality averages by period in Ise Bay and Mikawa Bay (1980-1989 and 2010-2019). **a)** N concentration distribution in 1980s. **b)** N concentration distribution in 2010s. **c)** P concentration distribution in 1980s. **d)** P concentration distribution in 2010s. **e)** Chl-a concentration distribution in 1980s. **f)** Chl-a concentration distribution in 2010s.

**5. Temporal Trends in Red and Blue Tide Events in Ise Bay and Mikawa Bay and Mitigation Strategies for Blue Tides in Aburagafuchi-Lake. (Figure S6)**

As shown in Figure **S6a**, the number of red tide occurrences has been significantly reduced since 1990 in Ise Bay and Mikawa Bay due to purification measures. However, despite the ongoing purification measures, the frequency of red tide occurrences has remained flat since 1995, suggesting that the effect of red tide countermeasures is limited due to the decline in nutrient concentrations. On the other hand, the decrease in the number of red tide occurrences has also reduced the frequency of mass deaths of marine organisms, and as a result, the number of blue tide occurrences has also decreased. Blue tides steadily decreased until around 1998, but their frequency has not decreased since then. Figure **S6b** shows that the oxygen status of the bottom sediments can be improved by covering the bottom of Aburagafuchi-lake with sand (Figure **S2b**). Therefore, we attempted to further suppress the occurrence of blue tides by covering part of the seabed with sand, but the effect of mitigating seabed hypoxia was limited because the target sea area is vast compared to Aburagafuchi-lake and additional measures such as removing accumulated sludge by dredging were required.

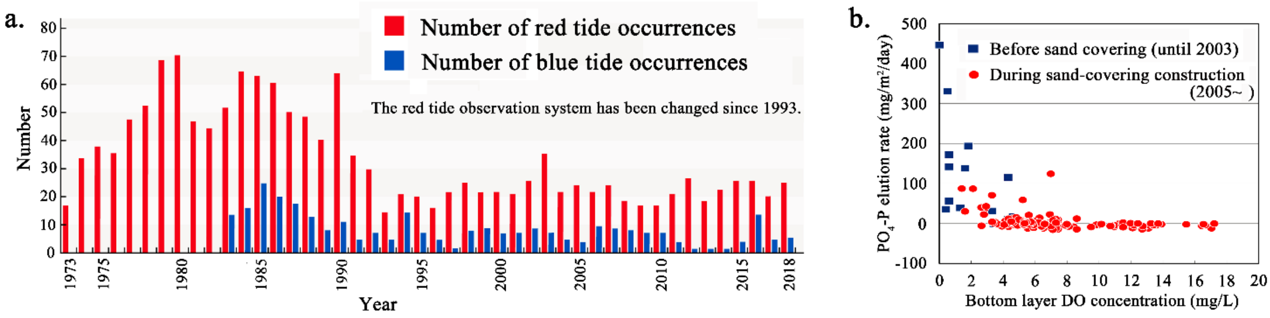


Figure **S6**. **a.** Changes in the annual number of red tides and blue tides occurring in Mikawa Bay and Ise Bay. **b.** The impact of covering the lake bottom with sand after dredging in Aburagafuchi-lake on bottom oxygen levels (DO) and inorganic phosphorus (Total P).

#### 6. Annual Changes in Catch Volume in Ise Bay and Mikawa Bay (mainly Mikawa Bay). (Figure S7)

Figures **S7a** and **S7b** show that the catches of benthic organisms such as mantis shrimp, conger eels, shrimp, and clams have decreased significantly. In particular, mantis shrimp, conger eels, and shrimp, which are benthic organisms that mainly feed on phytoplankton, have been on a downward trend since around 2011. In addition, the tiger pufferfish, a higher-order consumer that does not move much outside the bay, has also decreased significantly during the same period. Figure **S7c** shows that the catches of tiger pufferfish have also decreased despite the appearance of dominant year fish. On the other hand, the decrease in the catches of short-neck clams, which are benthic organisms with a narrow activity range, began around 1991, even earlier than these organisms. To prevent this decrease in catches, measures such as transplanting and releasing juvenile clams have been taken, but the catches of short-neck clams have continued to decrease significantly. Although these measures led to a temporary recovery after 2000, the catches have started to decrease again since 2012. This suggests that the decrease in nutrients is affecting phytoplankton. In Ise Bay and Mikawa Bay, it has been pointed out that the decline in phytoplankton may be disrupting the food chain, including zooplankton. Consequently, the degree of oligotrophy within the bay is hypothesized to be estimable through the concentration of Chl-a, a proxy for phytoplankton biomass. Therefore, it is thought that the level of oligotrophy in the bay can be estimated from the

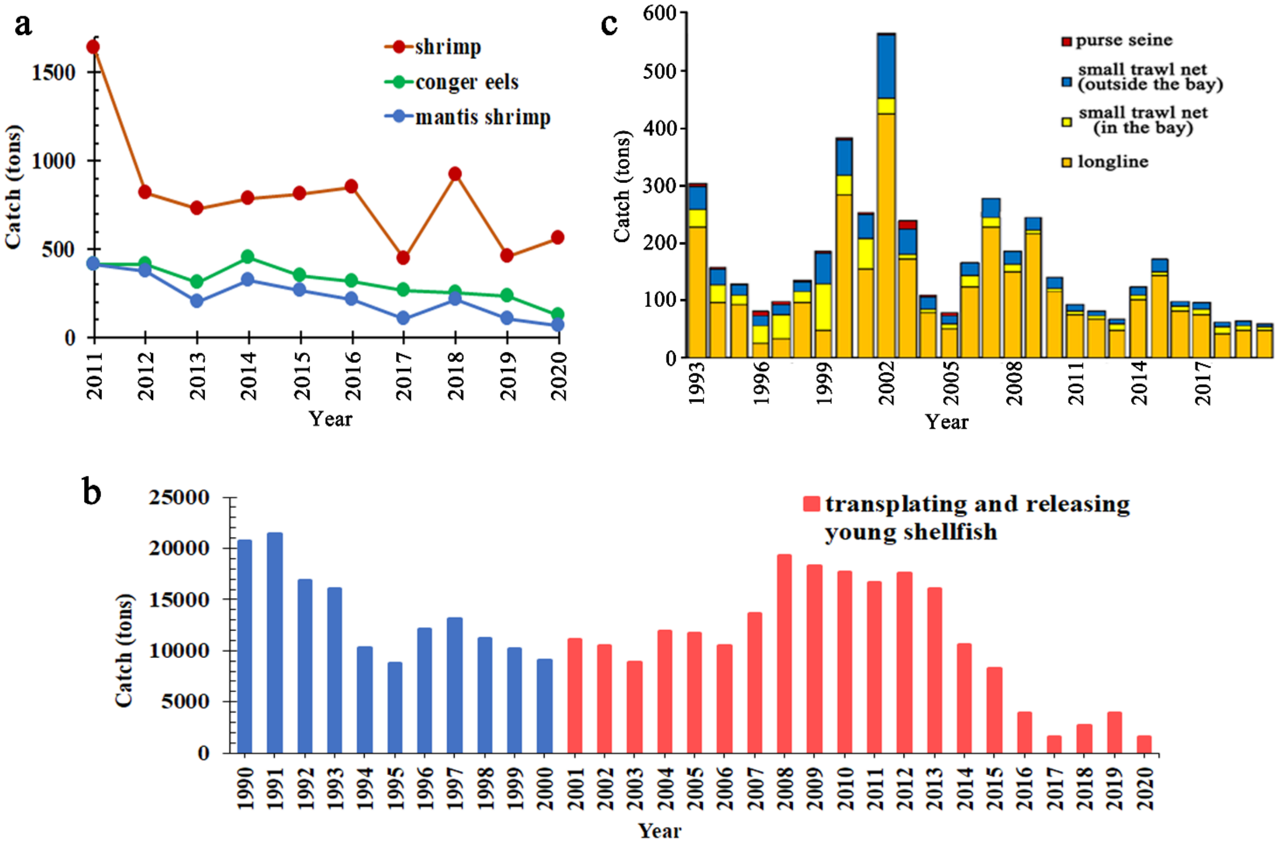


Figure **S7.** Trends in annual fishing catches in Mikawa Bay and Ise Bay. **a)** Catch volume of mantis shrimp, conger eel, and shrimp. **b)** Catch volume of short-neck clams. **c)** Catch volume of tiger pufferfish.

Chl-a concentration, which indicates the amount of phytoplankton. Among the benthic organisms, the catch and growth rates of short-necked clams, which are believed to be particularly sensitive to oligotrophic conditions, serve as critical indicators for assessing the extent of oligotrophy in the bay. Long-term monitoring of catch data for short-necked clams could provide valuable insights into the environmental changes occurring within the bay.

#### 7. Impact of Chlorophyll a Concentration on Clam Growth in Mikawa Bay. (Figure S8)

Figure **S8a** indicates that when the chlorophyll a concentration reaches approximately 20 µg/L, the growth

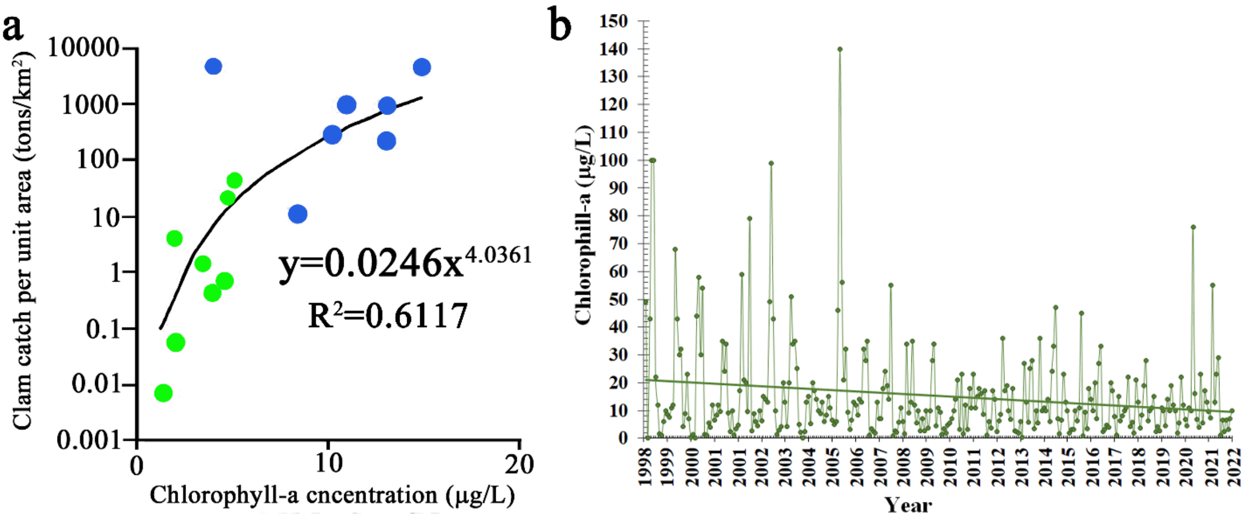


**Figure S8. a)** Relationship between short-neck clam yield per unit area and Chl-a content. Average catch volume at each fishing ground in the Seto Inland Sea from 2009 to 2011 (●); Average catch volume at each fishing ground in Mikawa Bay from 2009 to 2011 (●). **b)** Monthly Chl-a concentrations and their annual trends at environmental reference points K-4, K-5, K-6, and K-8 in Mikawa Bay (III).

rate of clams per unit area achieves saturation. Conversely, at a concentration of about 10 μg/L, clam growth becomes nearly negligible. Figure **S8b** further illustrates that the annual average chlorophyll a content at the environmental reference point in Mikawa Bay (III) declined from 20 μg/L to below 10 μg/L.

#### 8. Details of Aichi Prefecture's Social Experiment Conducted in Mikawa Bay. (Figure S9)

In this study, we examined how nutrient concentrations of nitrogen (N) and phosphorus (P) relate to Chlorophyll-a (Chl-a) levels according to the literature. Figure **S9a** reveals a weak correlation between N and Chl-a: while lower N levels slightly reduce Chl-a, the effect is minimal. Conversely, Figure **S9b** shows a strong positive correlation between P and Chl-a, where reduced P significantly decreases Chl-a levels. Thus, it appears that ocean oligotrophy is more strongly influenced by reductions in P than in N, due to P's greater impact on Chl-a.

Based on this assumption, efforts were made to elevate P concentrations in the effluent from wastewater treatment plants (WWTPs) discharging into Mikawa Bay. Figure **S9c** identifies two WWTPs involved in this study: the Yahagi-River WWTP (treatment volume: 277,560 $m^3$/day) and the Toyogawa WWTP (treatment volume: 91,879 $m^3$/day). From 2017 to 2021, P concentrations in these plants' effluent were intentionally increased for several months each year to counteract Mikawa Bay's declining P levels. Consequently, Figure **S9d** shows the P concentration in each WWTP's effluent rising from approximately 0.3 mg/L during normal operation to about 0.7 mg/L.

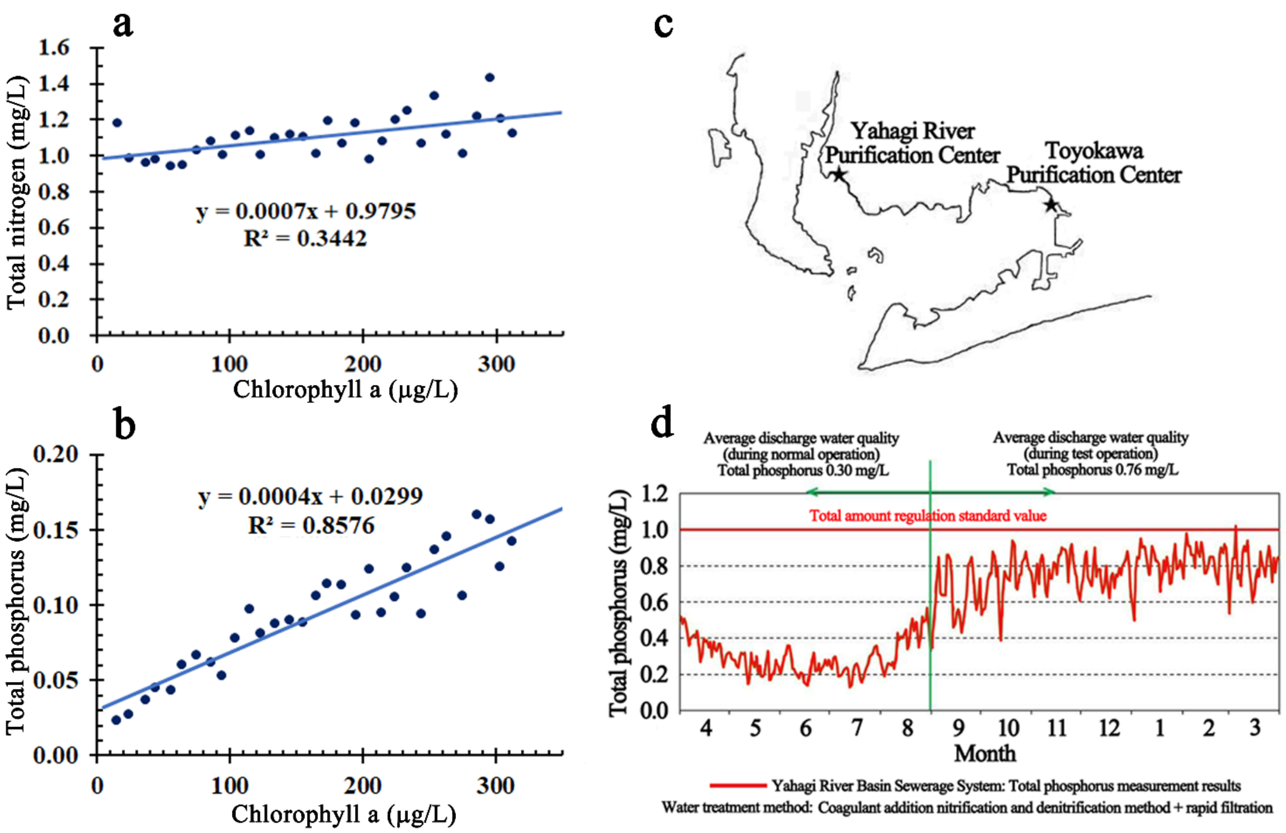


Figure **S9**. **a)** Correlation between total N and Chl-a. **b)** Correlation between total P and Chl-a. **c)** Location of the Yahagi-River and Toyogawa WWTPs. **d)** Changes in phosphorus concentration in treated water during normal and test operation at the Yahagi-River WWTP during the social experiment period.

#### 9. Rethinking Nutrient Dynamics and Chlorophyll-a Correlation Under Oligotrophic Conditions. (Figure S10)

As shown in Figure **S10a**, under oligotrophic conditions with low Chl-a concentrations, the correlation between P and Chl-a was evaluated after logarithmic transformation of the data, and the results showed that a power approximation was a better fit than a linear approximation. Similarly, the relationship between N and Chl-a was reevaluated, and it was found that there was little correlation with Chl-a when N was greater than 50 µg/L, and a negative correlation with Chl-a when N was less than 50 µg/L. This negative correlation can be explained by a power approximation, as shown in Figure **S10b**. To verify the reliability of these regression analyses, t-tests were performed. In this test, the values of the independent and dependent variables obtained as actual values were compared with the predicted values derived based on the regression analysis. A one-tailed test was used to evaluate whether the predicted values and the actual values were significantly different, and a paired two-sample t-test was performed, given that the data were paired data obtained from the same subject. The null hypothesis was set as "there is a significant difference between the predicted values and the actual values" and the significance level was set at less than 0.05.

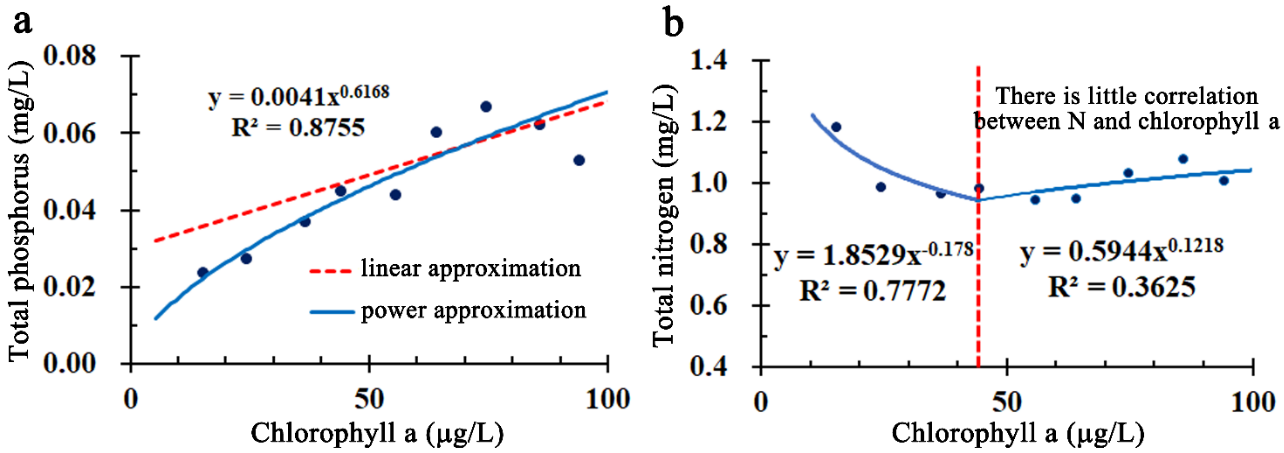


Figure **S10**. **a)** Correlation between total P concentration and Chl-a concentration under oligotrophic conditions. **b)** Correlation between total N concentration and Chl-a concentration under oligotrophic conditions.

#### 10. Phosphorus as the Key Driver for Phytoplankton Growth in Eutrophic Waters of Ise Bay and Mikawa Bay: A 25-Year Study. (Figure S11)

Figure **S11** illustrates the monthly relationships between P and Chl-a, N and Chl-a, and the N/P ratio and Chl-a at environmental reference points A-3, K-1, N-1, and K-2 in Ise Bay and Mikawa Bay from 1998 to 2023. At point A-3, the nitrogen concentrations are significantly higher and the nutrient status is better than at the other sites, due to the direct inflow of unregulated agricultural wastewater rich in nitrogen fertilizer that has not been absorbed by plants. Despite this, the Chl-a concentrations were lower than expected for a eutrophic sea area, possibly due to a deficiency in phosphorus. In contrast, points N-1 (Nagoya Port), K-1, and K-2 (Kinuura Port) receive treated water from Nagoya City and agricultural wastewater and septic tank effluent, respectively. These sites have high P and N values, but lower than those at A-3, with a more pronounced decrease in N than in P. However, Chl-a concentrations at these sites were similar to those at A-3, and there was an observed tendency for Chl-a values to increase when the N/P ratio was below 10. These results suggest that the increase in phosphorus concentration likely contributes to higher Chl-a concentrations in eutrophic waters, making phosphorus the main factor for phytoplankton growth.

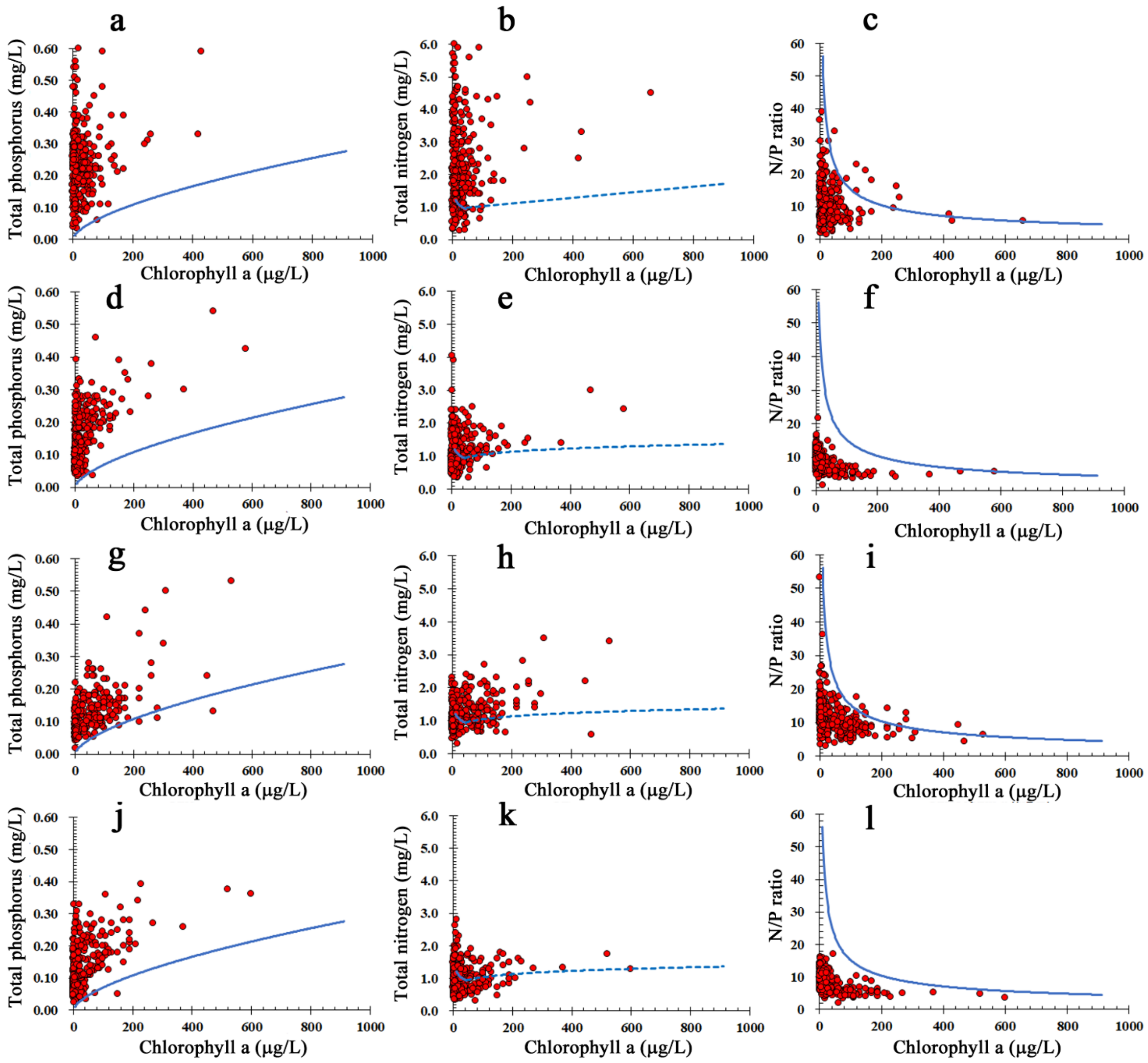


**Figure S11.** Monthly measurements of P, N, and Chl-a at the environmental reference points in Ise Bay and Mikawa Bay were plotted together with the respective response curves obtained by regression analysis of P, N, or N/P versus Chl-a. **a-c**) Environmental reference point A-3 (Kamino, Tahara City), where unregulated agricultural wastewater flows in. **d-f**) Environmental reference point K-1 in Kinuura Port. **g-i**) Environmental reference point N-1 in Nagoya Port. **j-l**) Environmental reference point K-2 in the southern part of Kinuura Port.

#### 11. Nitrogen Limitation as the Primary Factor for Phytoplankton Growth in Nutrient-Deficient Areas of Ise Bay and Mikawa Bay: A 25-Year Study. (Figure S12)

As shown in Figure **S12**, we compared the relationships between monthly P, N, or N/P and Chl-a measurements from 1998 to 2023 at Nagoya Port Environmental Reference Point N-4, Tokoname Offshore Environmental Reference Point N-5, and Ise Bay Environmental Reference Points N-6 and N-7, where nutrient deficiencies are relatively severe. We found that the P content was significantly lower at these points compared to others. This indicates that increasing N content leads to a higher N/P ratio and an increased Chl-a content, which aligns with the Chl-a value on the response curve when considering the N/P ratio as the independent variable. Therefore, we assume that the amount of N is the main limiting factor for phytoplankton growth in these sea areas.

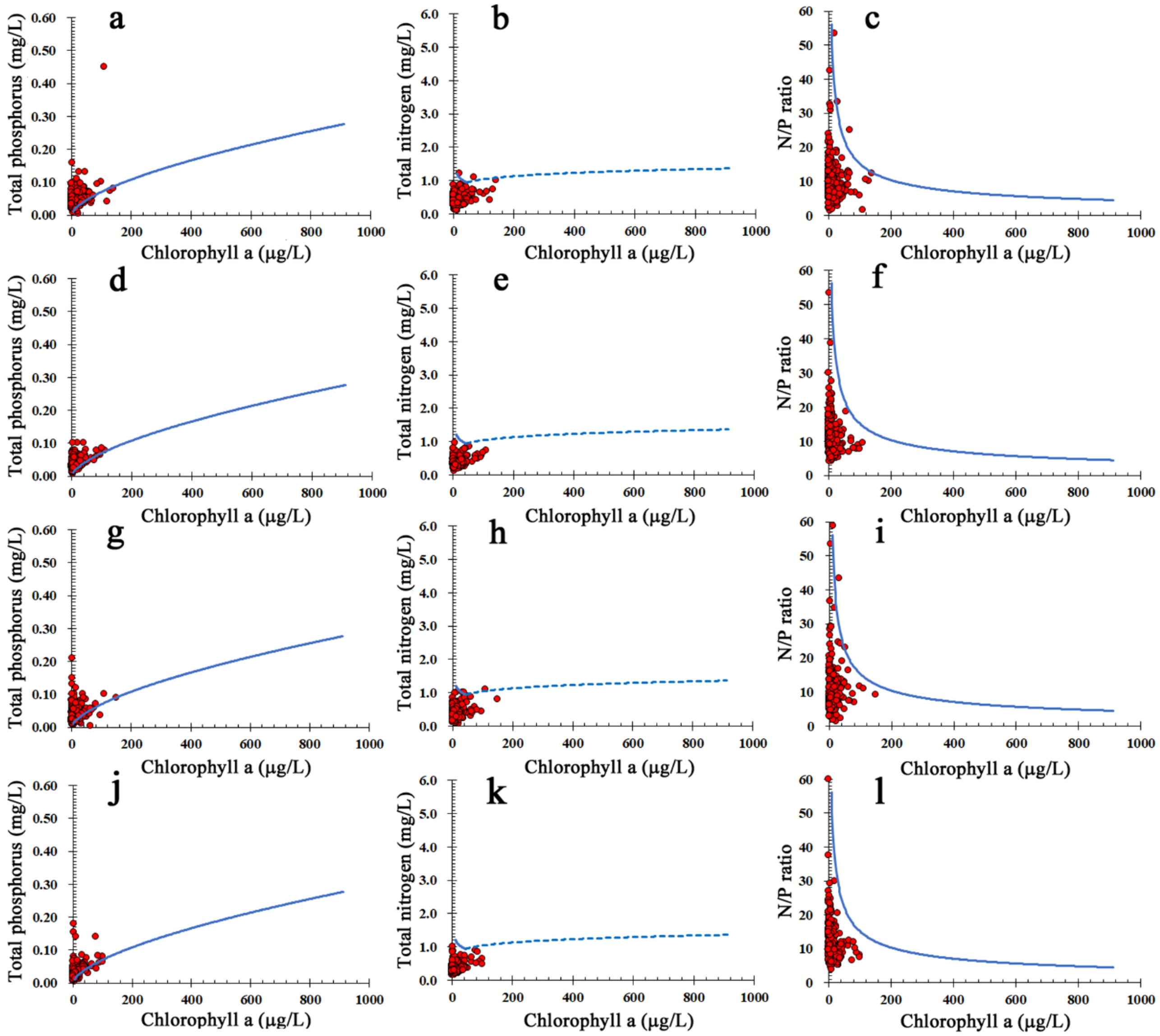


**Figure S12.** Monthly measurements of P, N, and Chl-a at the environmental reference points in Ise Bay and Mikawa Bay were plotted together with response curves obtained by regression analysis of P vs. Chl-a, N vs. Chl-a, and N/P vs. Chl-a. **a-c**) Environmental reference point N-4 (Nagoya Port) where treated sewage water from Nagoya City flows in. **d-f**) Environmental reference point N-5 (Tokoname offshore). **g-i**) Environmental reference point N-6 (Ise Bay). **j-l**) Environmental reference point N-7 (Ise Bay).

##### 12. Unexpected Chl-a Levels in Mikawa Bay: Discrepancies Between Nutrient Concentrations and Phytoplankton Growth. (Figure S13)

Figure **S13** shows the relationship between monthly P, N, or N/P ratio and Chl-a measurements from 1998 to 2023 at environmental reference points A-2, A-3, A-5, and A-13 in Mikawa Bay, where a social experiment is being conducted. Figures **S13a**, **S13b**, and **S13c** indicate that the concentrations of P and N are significantly higher at A-3 compared to other points in Mikawa Bay, yet the concentration of Chl-a is not as high as expected. For instance, Figures **S13j**, **S13k**, and **S13l** show that while the concentrations of N and P in A-2 are lower than those in A-3, Chl-a levels are almost the same as in A-3. This suggests that high Chl-a levels can occur without necessarily having high N and P concentrations.

The results indicate that Chl-a is maximized when the N/P ratio aligns with the response curve showing the power approximation of Chl-a and N/P ratio. When the N/P ratio deviates from this curve, the measured Chl-a is lower than the Chl-a predicted by the response curve, even if N and P values are high. Figures **S12f**, **S12i**, and **S12l** reveal that, based on Chl-a levels, Mikawa Bay is in an oligotrophic state unsuitable for phytoplankton growth, with nitrogen deficiency being the main limiting factor. This oligotrophic condition was also observed at all other environmental reference points in Mikawa Bay and at many points in Ise Bay.

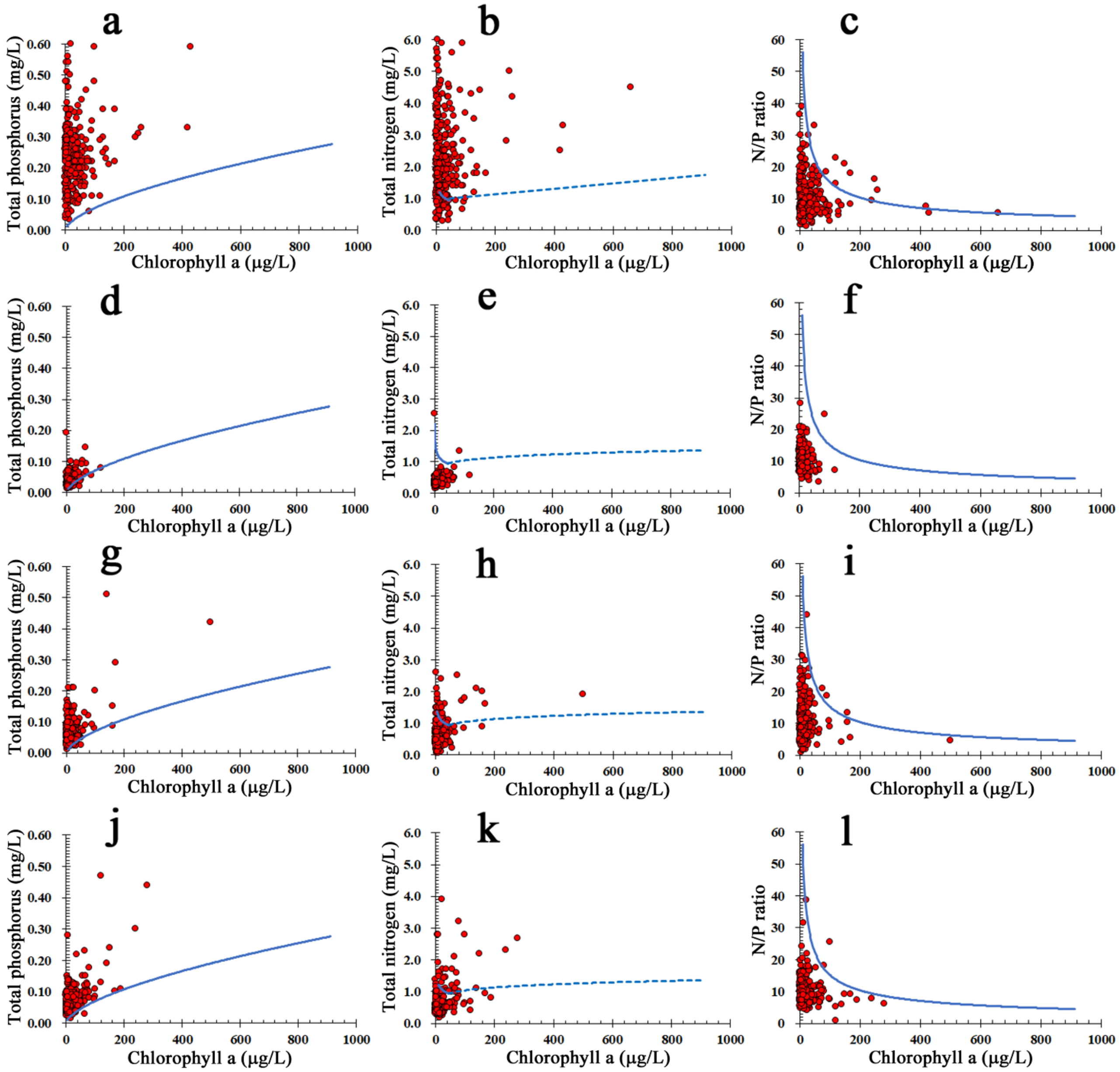


**Figure S13.** Monthly measurements of P, N, and Chl-a at environmental quality control points were plotted with response curves obtained by regression analysis of P vs. Chl-a, N vs. Chl-a, and N/P vs. Chl-a. a-c) Environmental reference point A-3 (Kamino, Tahara City), which receives unregulated agricultural wastewater. **a-c**) Environmental standard point A-3 (Kamino, Tahara City), where unregulated agricultural wastewater flows in. **d-f**) Environmental reference point A-5 in the center of Mikawa Bay. **g-i**) Environmental reference point A-13, adjacent to the Toyogawa WWTP. **j-l**) Environmental reference point N-7 (Ise Bay).

#### 13. Iron Ions in the Surface Water of the Rivers Flowing into Mikawa Bay may be a Potential Source of Metal Ions for Chemodenitrification. (Figure S14)

Recent research has shown that, in addition to biological denitrification, abiotic nitrate reduction (chemical denitrification) by minerals and organic matter in soil water is also significant. Particularly, the catalytic action of $Fe^{2+}$ ions in chemical denitrification has garnered attention. When chemical denitrification occurs in enclosed seas or in connected rivers and lakes, it leads to a decrease in the amount of inorganic nitrogen in these enclosed bodies of water. Surface waters of these rivers and lakes contain higher concentrations of $Fe^{2+}$ and $Fe^{3+}$ ions compared to other metals, as illustrated in Figure **S14**.

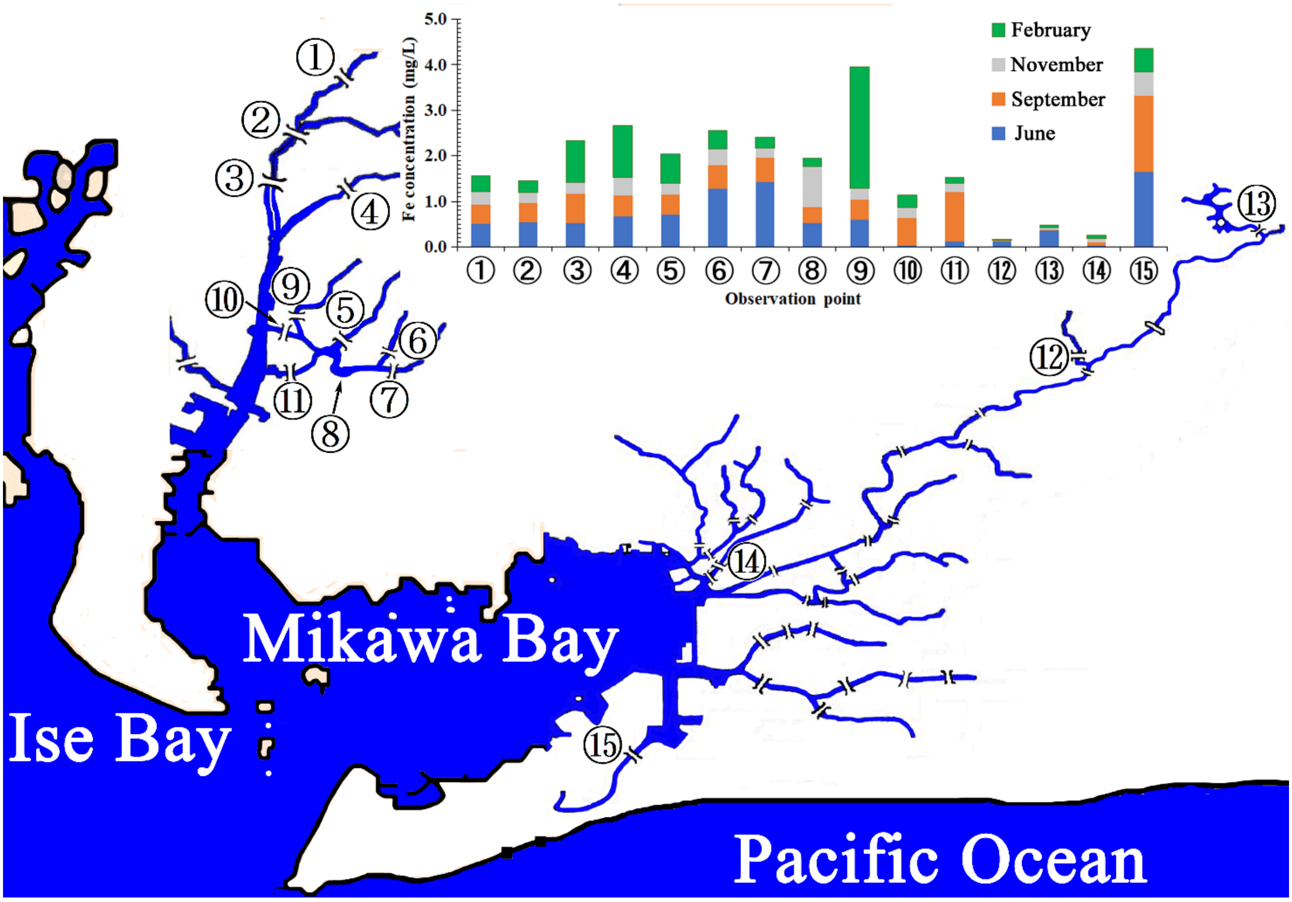


**Figure S14.** The iron content in water samples collected every three months at observation points ① to ⑮ in 2021 was measured using ICP-Mass. ①; Shinsakai Bridge, ②; Sakai River, ③; Ichihara Bridge, ④; Aizuma River, ⑤; Tansui Bridge, ⑥; Sakashita Bridge, ⑦; Sakashitakobashi Bridge, ⑧; Aburagafuchi, ⑨; Hieda Bridge, ⑩; Takahama Bridge, ⑪; Suimon Bridge, ⑫; Nagashino Bridge, ⑬; Hourai Bridge, ⑭; Yanagi Bridge, ⑮; Finakura Bridge.

#### 14. Anomalies in Chl-a Concentration and Assessment of Environmental Equilibrium in Ise Bay and Mikawa Bay. (Table S1).

A multiple regression analysis was conducted using observed values of nitrogen (N), phosphorus (P), and chlorophyll-a (Chl-a) at environmental reference points in Ise Bay and Mikawa Bay from 1998 to 2024. In this analysis, N and P served as explanatory variables, while Chl-a was the response variable, plotted within a three-dimensional coordinate system. To evaluate deviations from expected values, we calculated the number and rate of cases in which observed Chl-a values exceeded the response surface. These findings are summarized in Table **S1**. The analysis revealed that all outlier incidences were below 0.05. Specifically, for non-oligotrophic sites A-3 and N-1, the outlier incidences were 0.038 and 0.032, respectively. To estimate the 95% confidence intervals for the population outlier incidences, we applied the Z-score from the standard normal distribution (1.96) to the outlier incidences calculated from 312 samples. As a result, the confidence intervals were determined to be 0.060–0.017 for A-3 and 0.052–0.013 for N-1 (see Table **S1**). To ensure accuracy, we excluded sporadic outliers linked to environmental abnormalities such as red tides. These environmental anomalies were identified as data points with absolute errors greater than 50. However, evidence has not been established that relative errors greater than 50 are environmental anomalies. Following their removal, we reanalyzed the dataset to determine the number of outliers, outlier incidence, and mean absolute error (MAE), ultimately estimating the population outlier incidence. These revised results are detailed in Table **S1**.

**Table S1.** Rate of excessive outliers from the Chl-a response surfaces for monthly samples from Mikawa Bay and Ise Bay, and estimates of the expected rate of excessive Chl-a outliers in these Bays.

| Population | Sample Size | Outliers | | CI | | Anomalies* | Outliers-2** | | CI-2*** | | MAE |
|---|---|---|---|---|---|---|---|---|---|---|---|
| | | No. | Rate | Max. | Min. | No. | No. | Rate | Max. | Min. | |
| A-2 | 312 | 4 | 0.013 | 0.025 | 0.000 | 2 | 2 | 0.006 | 0.009 | 0.000 | 3.0 |
| A-3 | 312 | 12 | 0.038 | 0.060 | 0.017 | 8 | 4 | 0.013 | 0.025 | 0.000 | 14.9 |
| A-5 | 312 | 2 | 0.006 | 0.027 | 0.000 | 1 | 1 | 0.003 | 0.009 | 0.000 | 40.5 |
| A-13 | 312 | 4 | 0.013 | 0.016 | 0.000 | 0 | 4 | 0.013 | 0.025 | 0.000 | 18.8 |
| K-1 | 312 | 1 | 0.003 | 0.009 | 0.000 | 1 | 0 | 0.000 | 0.000 | 0.000 | 0.0 |
| K-2 | 312 | 1 | 0.003 | 0.009 | 0.000 | 1 | 0 | 0.000 | 0.000 | 0.000 | 0.0 |
| N-1 | 312 | 10 | 0.032 | 0.052 | 0.013 | 6 | 4 | 0.013 | 0.025 | 0.000 | 16.5 |
| N-4 | 312 | 4 | 0.013 | 0.025 | 0.000 | 0 | 4 | 0.013 | 0.025 | 0.000 | 3.7 |
| N-5 | 312 | 0 | 0.000 | 0.000 | 0.000 | 0 | 0 | 0.000 | 0.000 | 0.000 | 0.0 |
| N-6 | 312 | 4 | 0.013 | 0.025 | 0.000 | 1 | 3 | 0.010 | 0.020 | 0.000 | 4.3 |
| N-7 | 312 | 0 | 0.000 | 0.000 | 0.000 | 0 | 0 | 0.000 | 0.000 | 0.000 | 0.0 |

In this table, outliers refer only to observed values that exceed the theoretical values.

Abbreviation: MAE; Mean Absolute Erorr, CI; Confidence Interval

* Anomalies: Outliers with absolute error of 50 or more

** Outliers-2 with absolute error less than 50

***Confidence interval for population outlier rate of outlies-2

#### 15. Developed Views of the 3D Scatter Plots of A-3 and A13 in Figures 4a and 4d. (Figures S15 and S16)

Figures **S15a** and **S16a** show the observations (●) of samples collected monthly at the environmental reference points A-3 and A-13 from 1998 to 2023. These are plotted in a three-dimensional coordinate system with P and N as explanatory variables and Chl-a as the response variable, and the response surfaces modeled based on multiple regression analysis. Furthermore, Figures **S15b–e** and **S16b–e** show the projections of these 3D scatter plots and response surfaces from different angles.

Specifically: Figures **S15b** and **S16b**: Top views from the direction of 600 µg/L Chl-a, including the observed values within the response surface. Figures **S15c** and **S16c**: Bottom views from the direction where Chl-a is at 0µg/L. Figures **S15d** and **S16d**: Side views projected onto the plane consisting of the T-P and Chl-a axes. Figures **S15e** and **S16e**: Side views projected onto the plane consisting of the T-N and Chl-a axes.

From these projection views, it was confirmed that Figures **4a** and **4b** adequately represent the scatter plots of monthly observations and their corresponding response surfaces at environmental reference points A-3 and A-13. Note that in these figures, for clarity, the Chl-a axis has been truncated to include more than 99% of the Chl-

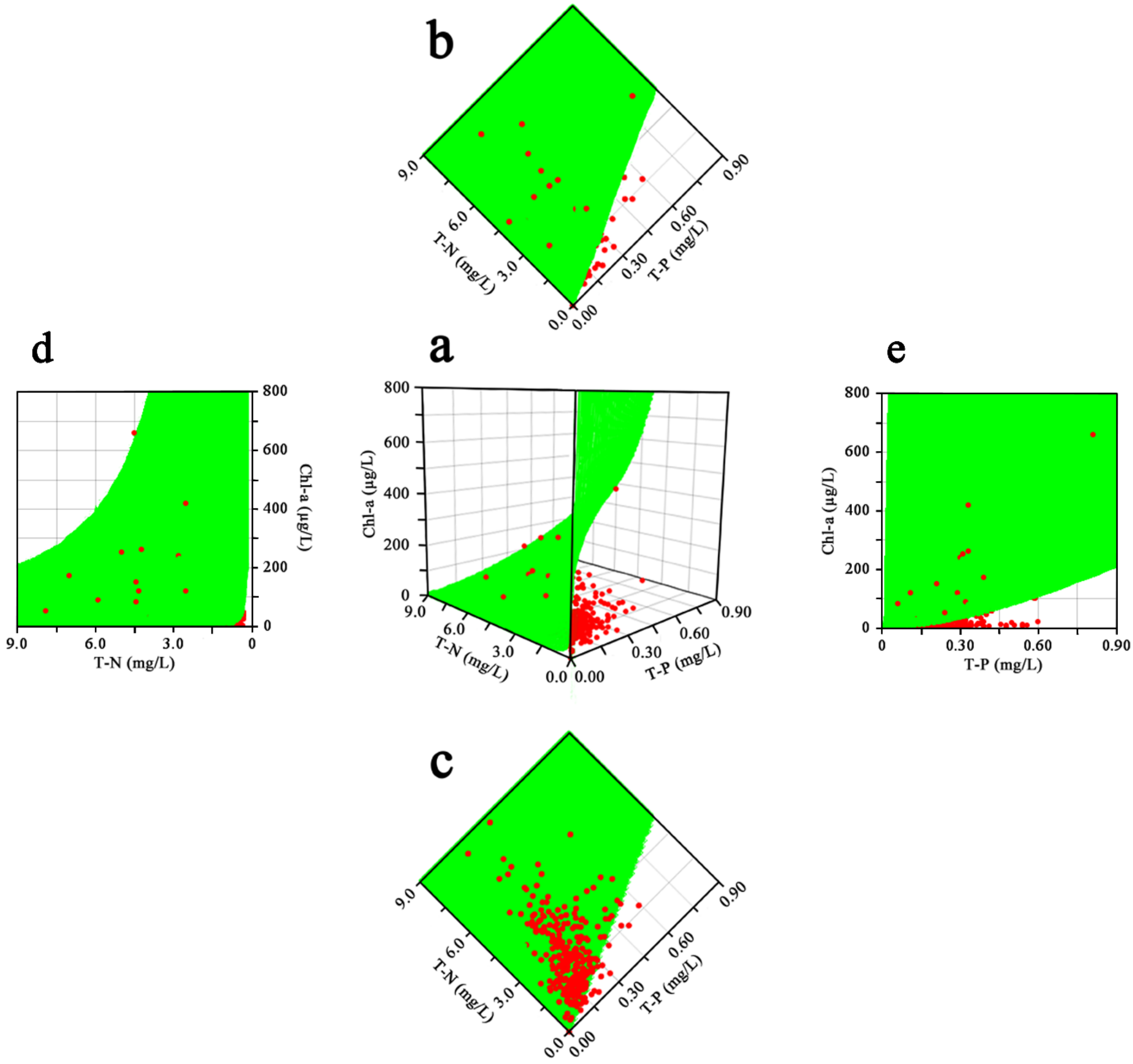


Figure **S15**. **a**) Scatter plot of the observed values (●) of samples taken monthly at environmental reference point A-3, and a response surface based on multiple regression analysis in a three-dimensional coordinate system with P and N as explanatory variables and Chl-a as the target variable. **b**) Its top projection. **c**) Its bottom projection. **d**) Its side projection on the T-N and Chl-a sides. **e**) Its side projection on the T-P and Chl-a sides.

a values, and some frames have been omitted.

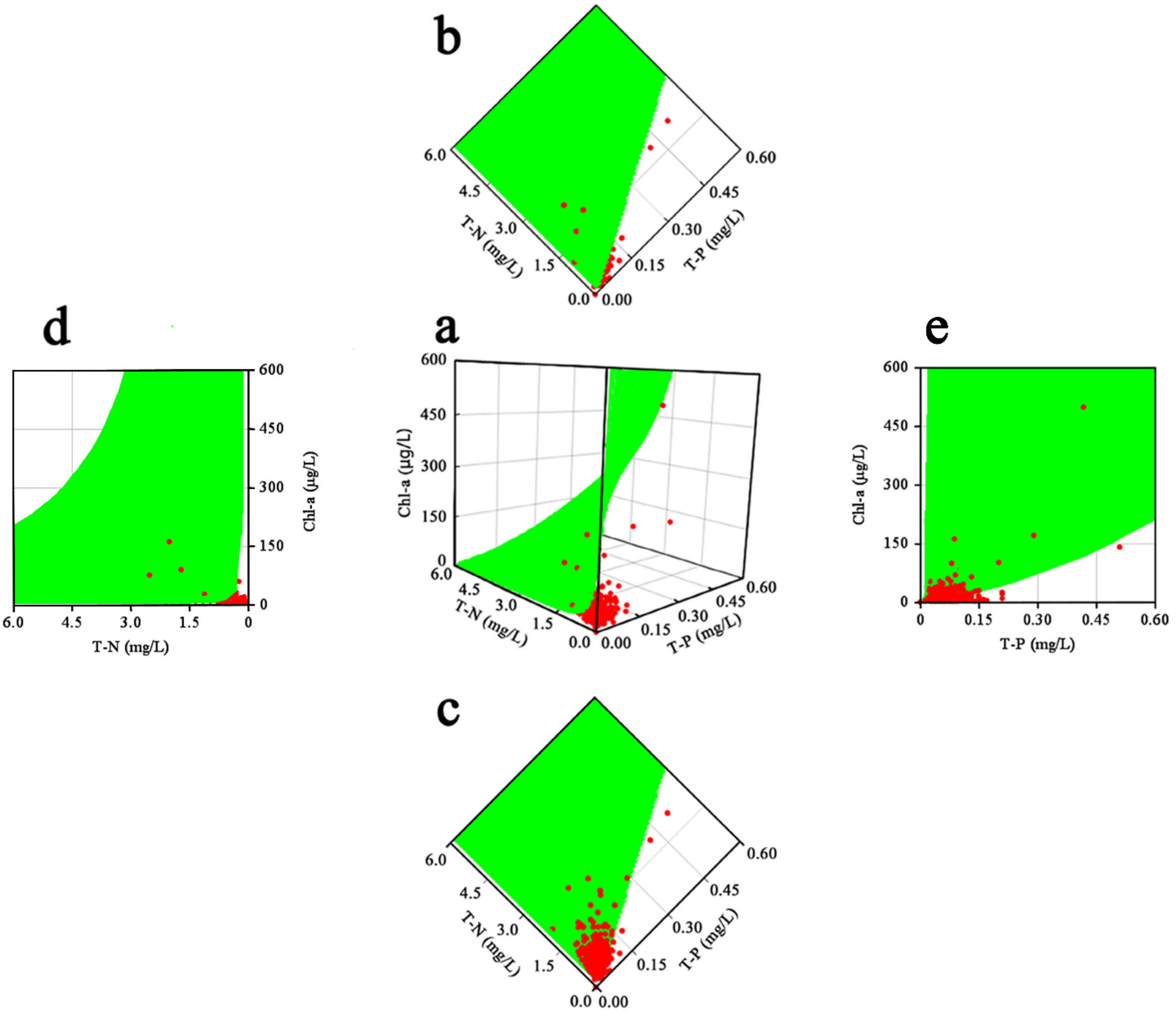


Figure **S16**. **a**) Scatter plot of the observed values (●) of samples taken monthly at environmental reference point A-13, and a response surface based on multiple regression analysis in a three-dimensional coordinate system with P and N as explanatory variables and Chl-a as the target variable. **b**) Its top projection. **c**) Its bottom projection. **d**) Its side projection on the T-N and Chl-a sides. **e**) Its side projection on the T-P and Chl-a sides.

#### 16. Investigating Nutrient Concentration Changes near the Wastewater Treatment Plant (WWTP) Outlet (Figure S17)

In Aichi Prefecture, a social experiment was conducted where treated water containing twice the normal amount of N and P was discharged from the Yahagi-River and the Toyogawa WWTPs over five months. This was done to investigate how the concentration distribution of these elements changed compared to their normal levels. The results are summarized in Figure **S17**.

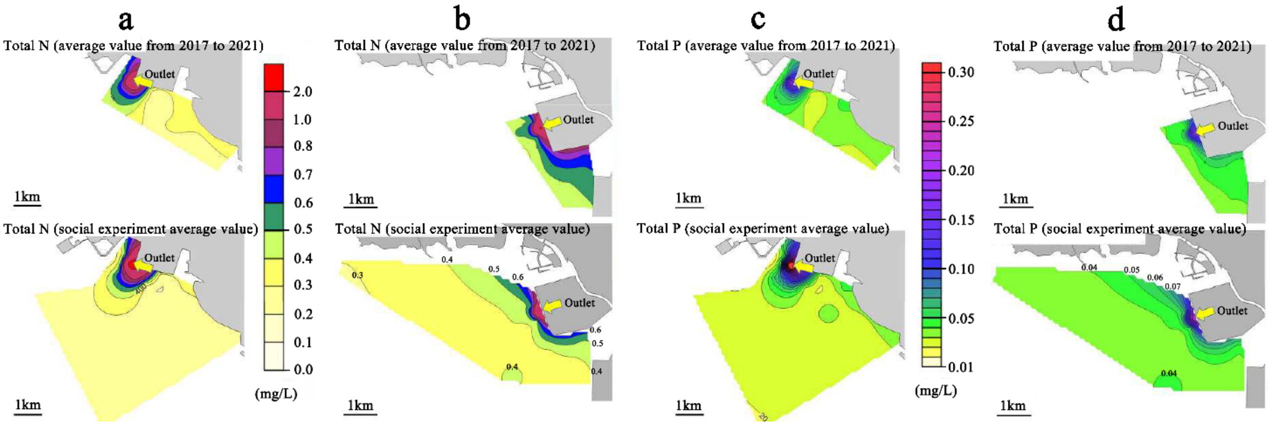


**Figure S17.** A comparison of N and P concentration distribution maps in water discharged into Mikawa Bay from the Yahagi-River WWTP and Toyogawa WWTP (November 2022 to March 2023) with concentration distribution maps in Mikawa Bay for the same period over the past five years (2017 to 2021). **a)** N distribution in effluent from the Yahagi-River WWTP. **b)** Distribution map of N in the bay in the discharge from the Toyogawa WWTP. **c)** Distribution map of P in the bay in the discharge from the Yahagi-River WWTP. **d)** Distribution map of P in the bay in the discharge from the Toyogawa WWTP.
